# Goal-dependent tuning of muscle spindle receptors during movement preparation

**DOI:** 10.1101/2020.03.06.981530

**Authors:** Stylianos Papaioannou, Michael Dimitriou

## Abstract

Voluntary movements are believed to be advantageously prepared before they are executed, but the neural mechanisms at work have been unclear. For example, there are no overt changes in skeletal muscle activity during movement preparation. Here, using a delayed-reach manual task, we demonstrate a decrease in the firing rate of human muscle afferents (primary spindles) when preparing stretch rather than shortening of the spindle-bearing muscle. This goal-dependent modulation of proprioceptors begun early after target onset but was markedly stronger at the latter parts of the preparatory period. In two additional experiments, whole-arm perturbations during reach preparation revealed a congruent modulation of stretch reflex gains of shoulder and upper arm muscles. Our study shows that movement preparation can involve sensory elements of the peripheral nervous system. We suggest that central preparatory activity can also reflect sensory control, and preparatory tuning of muscle spindle mechanoreceptors is a component of planned reaching movements.

## Introduction

A key mission in sensorimotor neuroscience is to understand the function and consequence of “preparatory activity”, that is, the vigorous changes in neural activity that occur in multiple areas of the brain before onset of a voluntary reaching movement. Although the firing of such ‘preparatory’ neurons located in e.g., the premotor cortex has been linked to a variety of factors such as movement direction/extent^1,2^ and visual target location^3^, the specific function of preparatory activity has remained unclear. A previous claim that preparatory activity represents a subthreshold version of movement-related cortical activity^4^ has been contradicted more recently in support of the notion that preparation sets another initial dynamical state that promotes execution of the planned movement^5,6^. But it is unclear what this initial state actually entails and by which neural mechanisms exactly the benefits of movement preparation are realized. For example, although preparation benefits performance by lowering reaction time^7-9^, with longer preparation delays generally leading to better movement quality^10^, there are no overt changes in skeletal muscle activity during movement preparation. Moreover, recent behavioral findings indicate that movement preparation is mechanistically independent from movement initiation, with a distinct neural basis^11^.

Little attention has been placed thus far on the possibility that preparatory activity may also reflect control of sensory (i.e., proprioceptive) elements located in the peripheral nervous system. The aim of the current study was to investigate the impact of goal-directed movement preparation on muscle spindle output and assess any implications for ‘reflex’ motor responses. Independent modulation of spindle sensitivity/gain to dynamic muscle stretch (via the γ motor or ‘fusimotor’ system) could function as movement-related preparation that does not determine concurrent skeletal muscle activity, but can nevertheless affect the execution of movement through influencing stretch reflex responses of all latencies i.e., both Short Latency Reflex (SLR) responses engaging spinal circuits and Long-Latency Reflex (LLR) responses involving supraspinal centers. In other words, we hypothesize that preparatory activity in the brain may also underlie goal-appropriate patterns of antagonist engagement, by selectively modulating spindle output and negative feedback (i.e., mechanical resistance) of stretching muscles during planned reaching movements.

In what follows we describe positive findings generated by three independent but complementary experiments, each employing a different group of human participants. One experiment focused on recording muscle afferent activity from hand and digit actuators using microneurography (experiment ‘1’), and the other two experiments utilized a robotic manipulandum platform to study ‘reflex’ motor responses at the level of the whole arm (experiments ‘2’ & ‘3’). To our knowledge, experiment ‘1’ represents the first instance where muscle afferent activity was recorded in a context involving both a dedicated movement preparation period and active reaching. Recording from single muscle afferents rather than single fusimotor efferents is feasible but also preferable in our paradigm involving fully-alert active humans. That is, the result of any substantial change in γ activity is a change in the output of the muscle spindle, and the spindle organ acts as an integrator of input from multiple fusimotor fibers^12^.

## Results

In experiment ‘1’, participants performed a classic center-out reaching task with the right hand while we simultaneously recorded hand kinematics, relevant electromyography (EMG) signals and single afferent activity from wrist or digit extensor muscles (Fig. 1a). Hand movements controlled the 2D position of a cursor on a monitor, and the participant’s task was to move the cursor to reach one of eight peripheral visual targets. The targets/trials were presented in a block-randomized manner, hence there was no systematic difference in kinematic history across a particular group of targets. On each trial, a target would suddenly turn into a red filled circle, representing the target ‘cue’, and participants were instructed to move to this target as soon as the ‘Go’ cue appeared (target turned into a green outline). The participants could ‘fail’ a trial if they were too late in reaching the target (see Methods for more details). Recording from single afferents during naturalistic active movement is very challenging due to the high incidence of electrode dislocations, and noise in the afferent signal also increases with muscle tension. Therefore, the delayed-reach task was designed to be short and compact, concentrating on one albeit fundamental experimental manipulation: visual target location. Figure 1b-c presents exemplary single-trial data pertaining to the same primary spindle afferent (type ‘Ia’ afferent). Despite no overt changes in kinematic variables or EMG during movement preparation, there was a decrease in the afferent’s firing rate when preparing to reach a target that required stretch of the spindle-bearing muscle (Fig. 1b). However, no such decrease occurred when preparing to move in the opposite direction that required shortening the muscle (Fig. 1c).

**Fig. 1:**
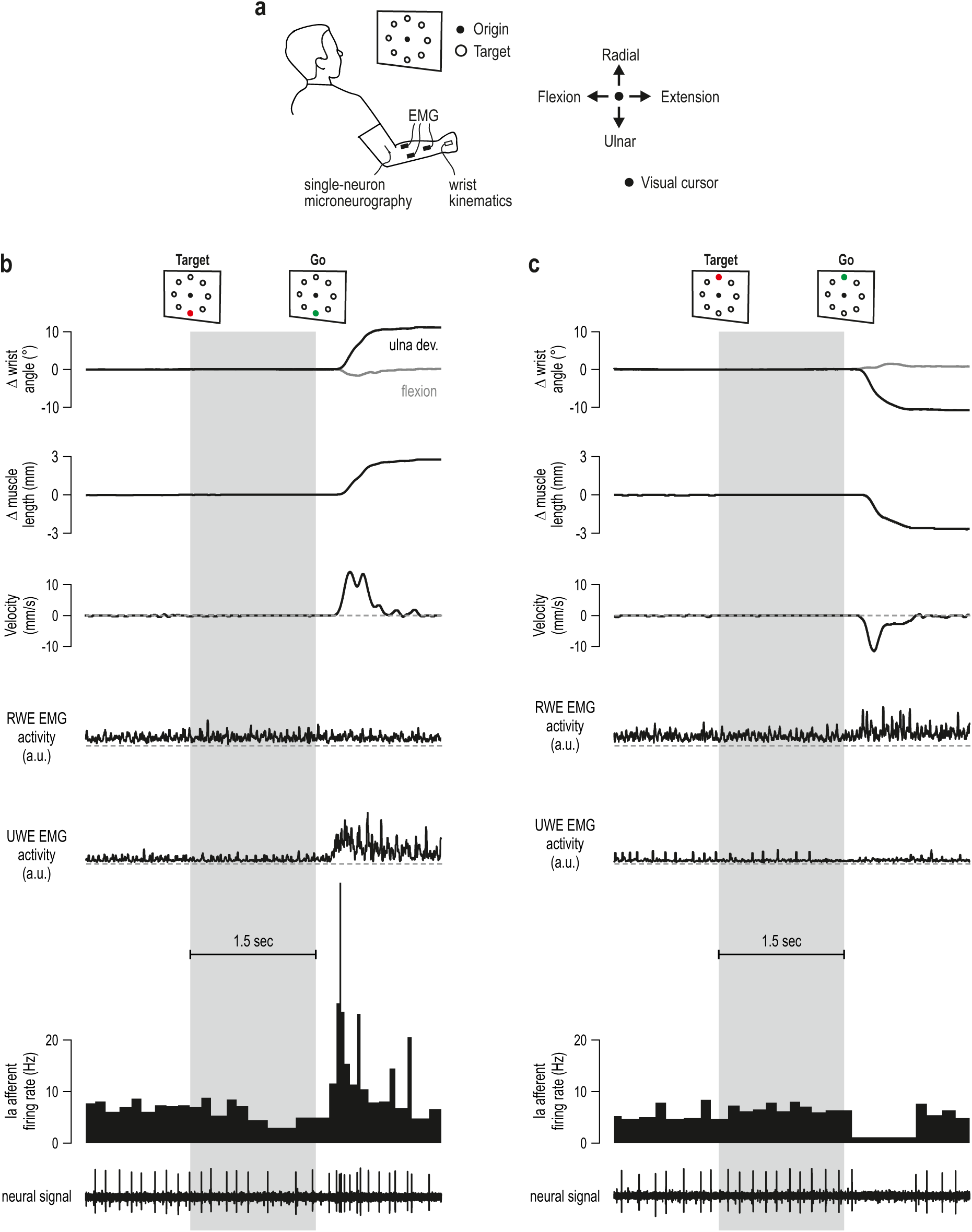
First experimental setup and representative single trial data. **a** The general setup of experiment ‘1’. Participants were asked to perform the classic delayed-reach task using their right hand. From an initial semi-pronated position, wrist flexion-extension moved a visual cursor in the horizontal dimension and wrist ulna-radial deviation moved the cursor in the vertical dimension. **b** Representative data from a single trial where reaching the target required ulna deviation of the wrist. Muscle length and velocity estimates pertain to the spindle-bearing muscle, which in this case is the Radial Wrist Extensor (RWE; i.e., *extensor carpi radialis*). Also shown is surface EMG from the Ulna Wrist Extensor muscle (UWE i.e., *extensor carpi ulnaris*) which mostly powered the reaching movement. Despite no overt changes in kinematics or EMG during the preparatory period (grey background), primary spindle afferent (‘Ia’) responses decreased, particularly at the latter half of this period. **c** The same neuron as ‘b’ but here the target was in the opposite direction, requiring radial deviation at the wrist and therefore shortening of the RWE. No decrease in firing rate was observed during the preparatory period. Throughout, dashed grey lines represent zero values.

From each participant in experiment ‘1’, we recorded muscle afferent activity from one of three muscles: the radial wrist extensor (‘*extensor carpi radialis*’), the ulna wrist extensor (*extensor carpi ulnaris*) or the common digit extensor (*extensor digitorum communis*). Single trials were categorized according to whether reaching the cued target required a substantial stretch or shortening of the spindle-bearing muscle (Fig. 2a). Despite no overt movement during the ‘preparatory period’ i.e., the period between onset of the target cue and onset of the ‘Go’ cue (Fig. 2b; top panel), type Ia population responses decreased when preparing to reach targets associated with stretch of the spindle-bearing muscle, relative to baseline (the latter is defined as values in the 0.5 sec epoch prior to target onset). The suppression effect appeared ∼80 ms after onset of the target cue and generally seemed to intensify closer to the onset of the ‘Go’ cue (Fig. 2b). This firing pattern could be seen at the level of single afferent spike-trains (Fig. 1b) and in all 8 recorded type Ia afferents (Fig. 2c), including those from digit extensor muscles that also stretch by wrist flexion. Single-sample t-tests confirmed the range of confidence intervals plotted in Figure 2c. Type Ia firing rates in all three epochs pertaining to subsequent muscle stretch (purple) were significantly different from baseline (epoch ‘1’: t(7)=-3.3, p=0.013; epoch ‘2’: t(7)=-3.1, p=0.017; epoch ‘3’: t(7)=-5.4, p=0.001), but this was not the case for targets associated with subsequent muscle shortening (‘blue’; all p>0.33). Indeed, a repeated measures ANOVA of the design 2 (target direction) x 3 (epoch) showed a main effect of target direction on Ia firing rates (‘stretch’ < ‘shortening’ targets) with F(1, 7)=10.8, p=0.013 and η _p_^2^=0.6, but the main effect and interaction involving the ‘epoch’ condition were not significant (p>0.1). However, complementary planned comparison tests confirmed the differences across confidence intervals displayed in Figure 2c. That is, when preparing to reach targets associated with muscle stretch, firing rates were lower in epoch ‘3’ vs. epoch ‘1’ with F(1, 7)=10.5 and p=0.014, but there was no significant difference in firing rate between epoch ‘2’ and ‘3’ (p=0.08). The results demonstrate that type Ia responses started to decrease early-on when preparing a movement that would stretch the spindle-bearing muscle, and this proprioceptive suppression intensified markedly closer to the onset of the ‘Go’ cue.

**Fig. 2:**
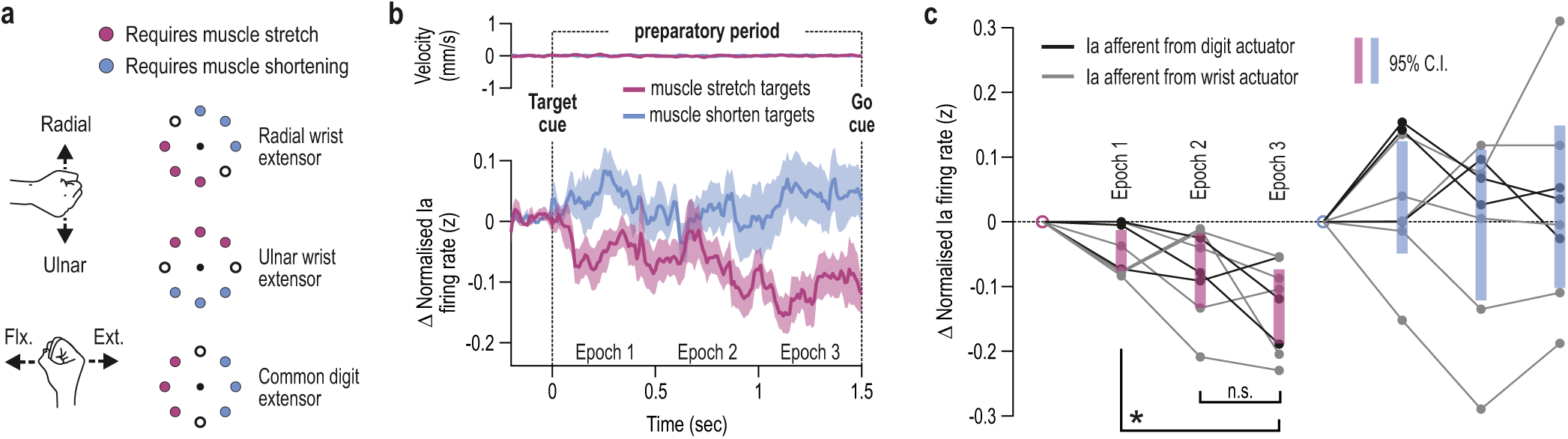
Goal-dependent tuning of muscle spindle receptors during movement preparation. **a** The visual targets were categorized based on whether reaching them required stretching or shortening of the spindle-bearing muscle. According to published physiological models for each muscle (see Methods), six targets represented clear and substantial change in muscle length, whereas two ‘intermediate’ targets (circle outlines) represented little or no muscle stretch or shortening. **b** Top panel represents mean stretch velocity of the recorded spindle-bearing muscles, essentially indicating no overt movement occurred in the preparatory period (see e.g., velocity scales in Fig. 1b-c and Supplementary Fig. 1). The bottom panel represents mean change in primary spindle afferent (‘Ia’) firing rates. All traces are aligned to onset of the target cue (time ‘0’). Purple and blue traces represent targets associated with stretch and shortening of the spindle-bearing muscle, respectively. Shading represents ±1 s.e.m. **c** Average Ia firing rates in the three epochs (‘1-3’) as shown in ‘b’. Thin grey lines represent individual Ia afferents from wrist extensor muscles and thin black lines represent Ia afferents from digit extensors. The shaded bars represent 95% confidence intervals and * p<0.05 following paired t-test. Same color scheme is used throughout. Goal-dependent decreases in tonic Ia firing rate may reflect a decrease in ‘dynamic’ fusimotor output to spindles; such fusimotor supply is known to have a much stronger effect on the spindles’ sensitivity to dynamic muscle stretch (i.e., gain).

As expected, kinematic and surface EMG signals showed no systematic variation in the preparatory period as a function of target cue. In certain cases, such as in the de-efferented spindles of the anaesthetized cat^13^, very small deviations in muscle length have been shown to impact spindle responses to stretch. Whether or not equivalently small deviations in hand kinematics could affect spindle responses in our paradigm, we had no reason to expect any systematic differences in kinematic variability during movement preparation with regard to the two groups of visual targets (Fig. 2a). Indeed, as reflected in Supplementary Figure 2, t-tests indicated no significant deviations from baseline during preparation for spindle-bearing muscle length (all p>0.36), velocity (all p>0.28), acceleration (all p>0.19) or EMG (all p>0.14), and no variable showed a trend or tendency towards the suppression pattern seen in spindle Ia responses prior to muscle stretch (i.e., purple epoch ‘3’ < purple epoch ‘1’ in Figure 2c).

We also recorded from four secondary spindle afferents (type ‘II’) and three Golgi tendon organ afferents (type ‘Ib’ encoding muscle-tendon tension) during the delayed-reach task. The same t-test analyses as above indicated no difference from baseline in type II firing rates (all p>0.36; Supplementary Fig. 3a), with no tendency towards the suppression pattern seen in spindle Ia responses. As all recorded type Ia afferents exhibited the goal-dependent suppression (Fig. 2c), and no consistent modulation was observed in type ‘II’ afferents, this suggests that a goal-dependent decrease in dynamic γ motor activity occurred when preparing to stretch the spindle-bearing muscle. These ‘dynamic’ fusimotor neurons only affect primary spindle receptors and, if everything else is equal (e.g., no movement), a substantive decrease in dynamic fusimotor output is known to induce some decrease in background (tonic) firing of type Ia afferents^14^ as shown in Fig. 2b-c. However, dynamic fusimotor supply has a much stronger effect on the gain of spindle output to dynamic muscle stretch^12,14^. A lowered spindle gain for stretching antagonists represents a degree of ‘on-line’ sensory attenuation of own action, and this proprioceptive suppression could also diminish counteractive negative feedback responses at all latencies. Interestingly, there also seemed to be a universal increase in type Ib firing rates regardless of target group (Supplementary Fig. 3b), and this did not parallel the state of the relevant parent EMG (Supplementary Fig. 3c). Although single-sample t-tests showed no difference from baseline in type Ib responses (all p>0.09), a 2 (target group) x 3 (epoch) repeated-measures ANOVA yielded a main effect of target group (F(1, 2)=21, p=0.044 and η_p_^2^=0.9), indicating that the increases in Ib firing rate were larger for shortening targets. There was no significant effect of epoch or interaction effect between target group and epoch (p>0.21). Because Golgi tendon organs and their respective Ib afferents are responsive to force produced by muscle fibers, this would appear to contradict the widely held belief of no changes in skeletal muscle activity during movement preparation; this universal increase in Golgi responses needs to be further confirmed by recording from a larger group of type Ib afferents. However, even if the type Ib effect is shown to be robust, it does not contradict the hypothesis that goal-dependent decreases in type Ia firing are due to suppression of γ neurons that are controlled independently of α motor neurons. Type Ib firing rates are unlikely to represent spindle state as intrafusal muscle fibers are known to make negligible contributions to muscle force^15^. Most important, however, the direction of the Ib effect (increase) is the same for both visual target groups (i.e., see purple and blue epoch ‘3’; Supplementary Fig. 3b), indicating that a different mechanism was at play here than the one controlling primary spindles. In other words, if the mechanism or underlying mechanical state responsible for modulating type Ib responses was also the one shaping type Ia responses, one would expect a larger suppression of type Ia activity for shortening targets, which was not the case (e.g., ‘blue’ in Fig. 2c). In addition, if the above were true, some systematic effect in type II afferent responses would also occur but none was found (Supplementary Fig. 3a).

Subsequent analyses also suggest an attenuation of dynamic γ influence on antagonist muscle spindles in delayed reach. As mentioned above, dynamic fusimotor activity has a weaker positive effect on tonic type Ia responses (‘offset’) and a much stronger positive effect on the spindle’s sensitivity to dynamic muscle stretch (‘gain’). Factors potentially shaping muscle spindle responses during active movement include spindle-bearing muscle length, velocity, acceleration and EMG, the latter used as a proxy for any ‘alpha-gamma co-activation’^16-18^. A forward stepwise regression indicated that only velocity and acceleration exerted a significant impact on type Ia population responses during reaching (Fig. 3a-b), with standardized beta coefficient=0.44 and p<10^−5^ for velocity, and beta=0.21 and p=0.025 for acceleration (R^2^=0.72, p<10^−5^). As expected, there was a significant relationship between velocity and empirical type Ia firing rates (Fig. 3d), with r=0.84 and p<10^−5^, but there was no significant one-to-one relationship between acceleration and firing rates (r=0.24, p=0.32). That is, spindles were unable to unambiguously ‘encode’ acceleration (Fig. 3f). Conducting equivalent analyses with type Ia firing rates from the radial wrist extensor muscle alone produced congruent results (Fig. 3c): only velocity and acceleration had a significant impact with beta=0.53 with p<10^−5^ for velocity, and beta=0.25 with p=0.01 for acceleration (R^2^=0.79, p<10^−5^). Similarly, the muscle’s Ia firing rate had a significant relationship with velocity but not with acceleration (Fig. 3e & 3g). The above findings contrast with those where hand reaching movements are performed without a preparatory delay. Specifically, in ‘unprepared’ manual reaching, the relative impact of acceleration on Ia responses from the radial wrist extensor is about twice as large (i.e., beta > 0.5) and greater than that of velocity, with a significant direct relationship existing between acceleration and Ia firing rate^17^.

**Fig. 3:**
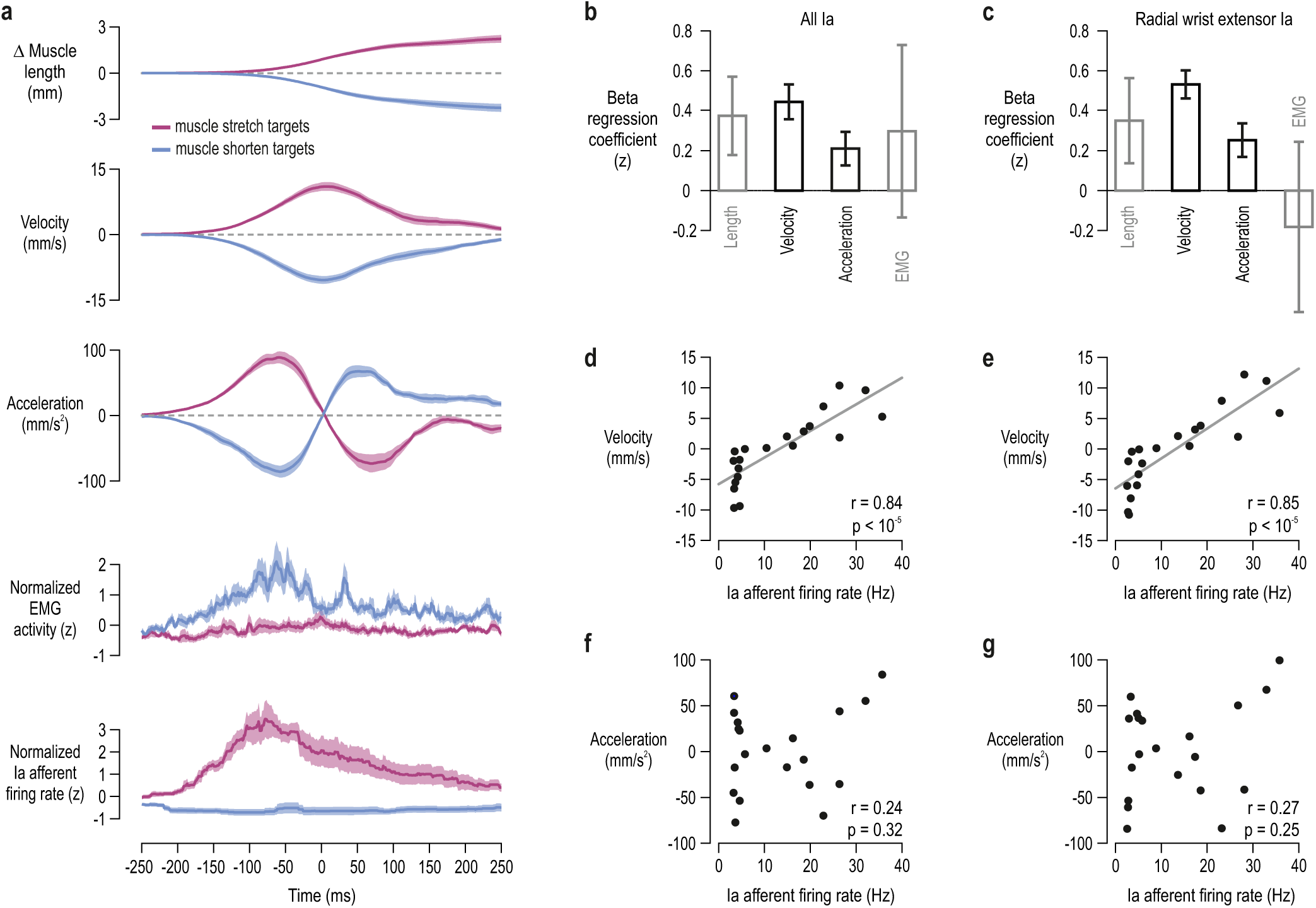
Muscle spindle receptors are relatively insensitive to acceleration during delayed reaching. **a** Type Ia population responses and associated spindle-bearing muscle kinematics and EMG. The signals were generated by averaging (mean) across the median responses of participants with whom a single afferent was recorded. Shading represents ±1 s.e.m. Signals are aligned to peak velocity (time ‘0’). **b** We used spindle-bearing muscle length, velocity, acceleration and EMG in a single regression as predictors of afferent firing rate (i.e., data shown in ‘a’, but down-sampled with a 50 ms moving average). ‘Beta’ regression coefficients are shown for facilitating comparison across predictors; these coefficients reflect the degree of change in the dependent variable (in units of s.d.) given a 1 s.d. change in the predictor variable. Error bars represent ±1 s.e.m. Black represents a statistically significant impact (p<0.05). **c** As ‘b’ but for spindle afferents originating from the Radial Wrist Extensor (RWE) muscle alone. Both velocity and acceleration were significant predictors but the impact of acceleration on Ia firing rate is ∼half of that observed when performing reaching movements in the absence of a preparatory period (beta > 0.5; see^17^). **d-e** As expected, there was a strong significant relationship between velocity and type Ia firing rates across all recorded Ia afferents (‘d’) and those of the RWE alone (‘e’). **f-g** There was no significant relationship between acceleration and firing rate across all recorded Ia afferents (‘f’), nor with those from the RWE (‘g’).

As mentioned above, changes in tonic Ia firing during movement preparation are presumed to be -at least partly-indicative of dynamic fusimotor activity levels. If the system sought to systematically attenuate fusimotor outflow to antagonist muscles, this implies that suppression of spindle output from stretching antagonists is beneficial for reaching performance. We show that a likely consequence of fusimotor attenuation is the decrease in spindle sensitivity to acceleration. But some sensitivity to acceleration clearly remained at the population level (Fig. 3b-c), in turn suggesting possible individual differences in the control of spindle sensors with implications for task performance. It is known that movement preparation benefits performance by lowering reaction time^7-9^, with a positive relationship existing between preparation delay length and movement quality^10^. Interestingly, although we found no relationship between type Ia firing rates observed during late preparation (i.e., epoch ‘3’) and reaction time (Fig. 4a-b), there was a strong relationship between wrist type Ia responses at epoch ‘3’ and time to peak velocity during reach, with r=0.9 and p=0.035. (Fig. 4c). Every unit increase in firing rate during preparation involved an additional 3 ms delay in reaching peak velocity; that is, the regression coefficient was 3. We found no equivalent relationship between this performance measure and kinematic variables or EMG (Supplementary Fig. 4). The relationship between time to peak velocity and Ia firing levels at late preparation seemed to extend beyond muscles that powered movement in the reaching task. For all but one afferent from digit extensors, the same relationship was found between type Ia firing rates and time to peak velocity (r=0.91, p=0.004, ‘b’ coefficient=0.301; Fig. 4d). Note that digit extensors can also affect execution of hand flexion via spinal and transcortical negative feedback circuits.

**Fig. 4:**
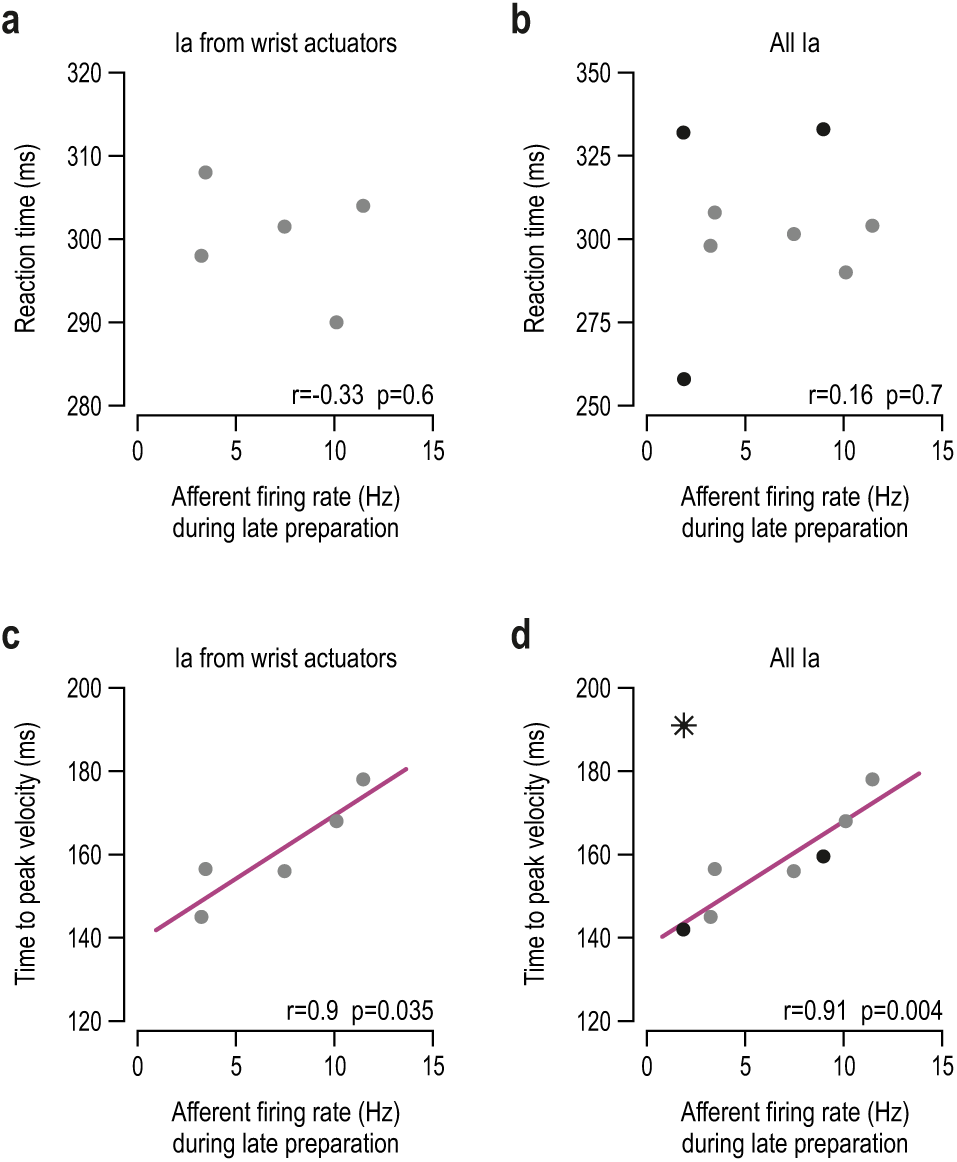
Spindle firing rates at late movement preparation predict performance during reaching. Throughout, each data point represents the average (median) value of a single participant/afferent across trials where reaching the target required stretch of the spindle-bearing muscle. The left column of panels pertains to wrist muscles (grey dots), and the right represents all Ia afferents, including those originating from digit extensor muscles (black). **a-b** The horizontal axes represent Ia firing rates during the late preparation epoch (i.e., epoch ‘3’ as defined in Figure 2b) and vertical axes represent reaction time i.e., the time between onset of the Go cue and onset of the reaching movement. **c** The vertical axes represent time between onset of reaching and the initial peak velocity of reaching movement; there was a strong positive relationship with tonic Ia firing from muscles engaged in powering hand movement in the current task (i.e., wrist actuators). **d** With the exception of one participant/afferent (black star), movement performance was well described by the same relationship (i.e., 3 ms delay in attaining peak velocity for every additional spike/sec). The relationship between spindle Ia responses at late preparation and subsequent reaching performance can be understood in terms of the spindle’s role in negative feedback circuits (i.e., stretch ‘reflexes’).

Indeed, muscle spindles play a central role in stretch reflex responses. A substantial goal-dependent suppression of spindle gains could lead to equivalent changes in negative feedback gains. In order to test this prediction of experiment ‘1’, we used a popular methodological approach for assessing stretch reflex function at the level of the whole upper limb. Namely, in experiment ‘2’, participants performed a version of the delayed-reach task by holding the graspable end of a robotic manipulandum with their right hand. They started each trial by bringing the hand at a central target (‘origin’; see Fig. 5a). The hand could then be slowly loaded in either the upper-left (135°) or lower-right direction (315°), or there could be no load. One of two possible targets would then be cued by turning red, and after either a ‘long’ or relatively ‘short’ preparatory delay (see Methods for more details) the hand would be perturbed in the same or opposite direction as the target i.e., either in the 135° or 315° direction (Fig. 5b). Importantly, even when perturbations were in the direction of the cued target, participants had to complete the planned movement themselves as the size of the perturbation was only about a third of the distance to the target. This ensured that movement control was required on every trial of this task, regardless of perturbation direction. Figure 5c-e displays the median responses of a representative participant. Despite identical displacement during the haptic perturbations, visual inspection of the EMG signal from the unloaded pectoralis indicates a clear difference at spinal SLR latencies as a function of cued target (i.e., 25-50 ms following perturbation onset; Fig. 5c). This difference is congruent with the afferent findings: a relative suppression of the SLR response when preparing to stretch the pectoralis (purple) rather than shorten it (blue). This suppression completely disappeared at high background activation levels of the pectoralis, induced by an external load applied prior to the haptic perturbation (Fig. 5e). A goal-dependent suppression of LLRs (i.e., EMG 75-100 ms post-perturbation) was evident across all load conditions.

**Fig. 5:**
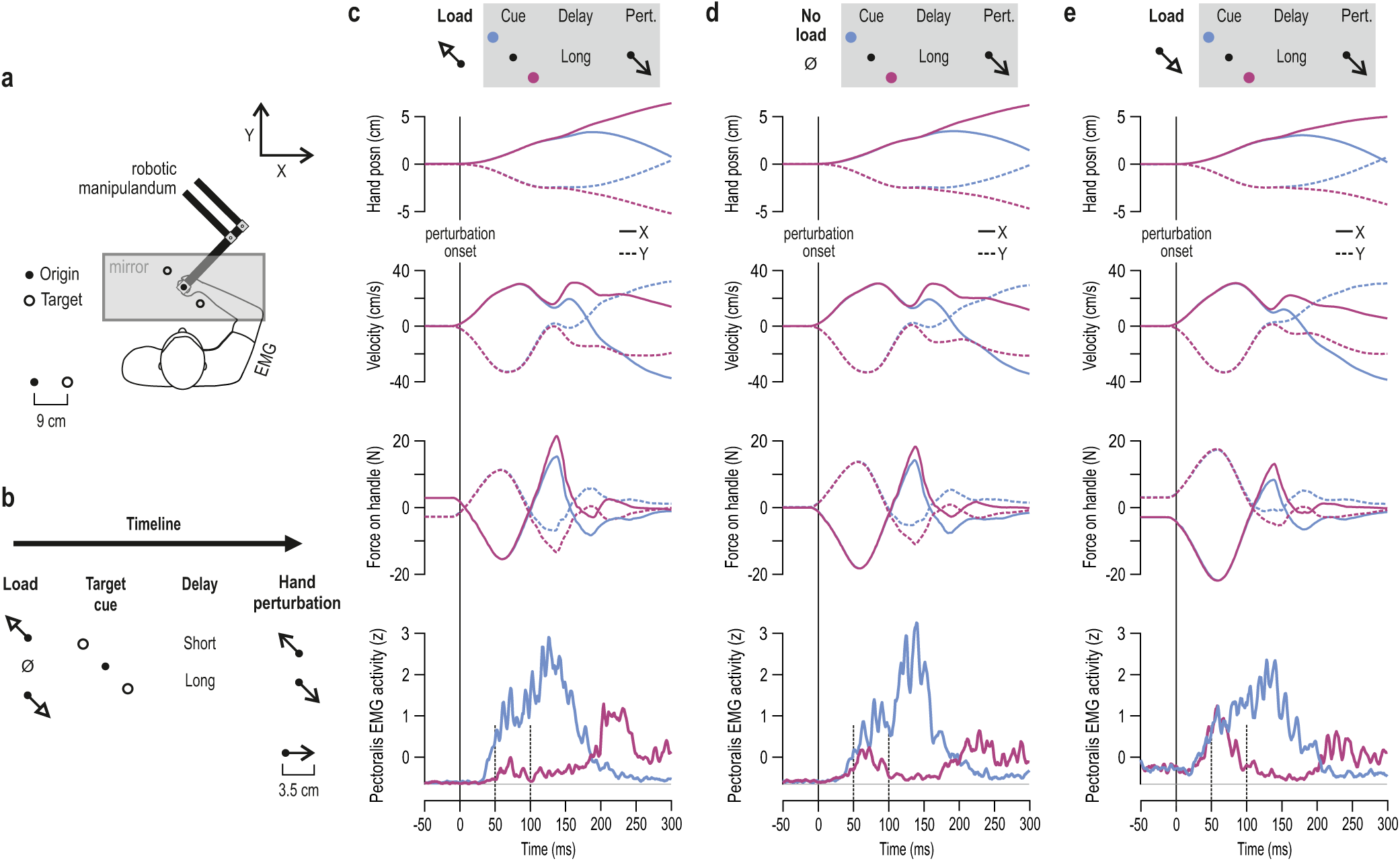
The second experiment and representative data from a single participant. **a** The general setup of experiment ‘2’. Participants held the graspable end of a roboric manipulandum. Vision was directed at a one-way mirror, on which the contents of a monitor were projected. Hand position was represented by a visual cursor. Although not shown here, the right forearm rested on an air-sled and the hand was immobile around the wrist (see Methods for more details). **b** The timeline of experimental manipulations. Each trial begun by slowly loading the hand in the upper left direction (i.e., -X and +Y direction) or lower right direction (+X and -Y), or there was no load (‘null’ load). The participants had to maintain the hand immobile at origin despite any loading. One of two visual targets was then suddenly cued (turned red) and this state lasted for a relatively short delay (250 ms) or long delay (750 or 1250 ms; see Methods). These preparatory delays correspond to the middle of epochs ‘1-3’ (Fig. 2b-c). At the end of the delay the hand was rapidly perturbed towards or in the opposite direction of the cued target. The perturbation lasted for 150 ms; at its end the ‘Go’ signal was given (cued target turned green) and movement to the target had to be completed. Cursor position was frozen during the perturbation. Trials were block-randomized, hence perturbation direction was unpredictable even after experiencing a particular load and cue. **c-e** Relevant median signals from a single participant when perturbations stretched the pectoralis muscle following a 750 ms preparatory delay, after first applying a load in the direction of pectoralis shortening(‘c’), when there was no external load (‘d’) and when applying a load in the direction of pectoralis stretch, promoting increased pectoralis activity for maintaining the start position (‘e’). Data are aligned to the onset of the position-controlled haptic perturbation (time ‘0’), defined as the point where movement speed reached 5% of initial peak value.

Figure 6a-c represents the equivalent to Figure 5c-e for all participants. The same trends can be seen in continuous EMG signals, that is, a goal-dependent suppression of pectoralis SLR and LLR. To concentrate on the effect of cued target while accounting for known effects, such as the universal increase in SLR magnitude that accompanies muscle loading^19-21^, the EMG signals for each muscle, load condition and delay condition were contrasted (subtracted) as a function of target cue. This effectively isolated any effect of target cue on SLR responses (but see also LLR analyses across loads). Note that in experiment ‘2’ and ‘3’ we only analyzed EMG signals from stretching muscles (i.e., particular pairs of muscle and perturbation direction) in order to concentrate on stretch reflex responses. When the preparation delay was long (Fig. 6d), single-sample t-test indicated a significant suppression of pectoralis SLR when preparing stretch in the ‘muscle unloaded’ condition, i.e., when an external (pre-)load was applied in the direction of muscle shortening (t(13)=-3.5, p=0.004). There was also a significant effect of goal in the no-load condition (t(13)=-2.5, p=0.025), but there was no relative suppression as a function of target cue in the ‘muscle loaded’ condition i.e., when the external load was applied in the direction of pectoralis stretch (t(13)=-0.23, p=0.82). Interestingly, when the preparation delay was relatively ‘short’ (250 ms; Fig. 6e-h), there was no suppression of SLRs when an external load was applied in either direction (p>0.8) but there was a relatively weak suppression effect in the no-load condition, with t(13)=-0.25, p=0.025. A congruent pattern of effects was observed for the posterior deltoid muscle (Supplementary Fig. 5). Specifically, when the preparation delay was long, there was a significant suppression of deltoid SLR in the ‘muscle unloaded’ condition (t(13)=-3.7, p=0.002). There was also significant suppression of SLR in the no-load condition when the delay was short (t(13)=-3.3, p=0.006).

**Fig. 6:**
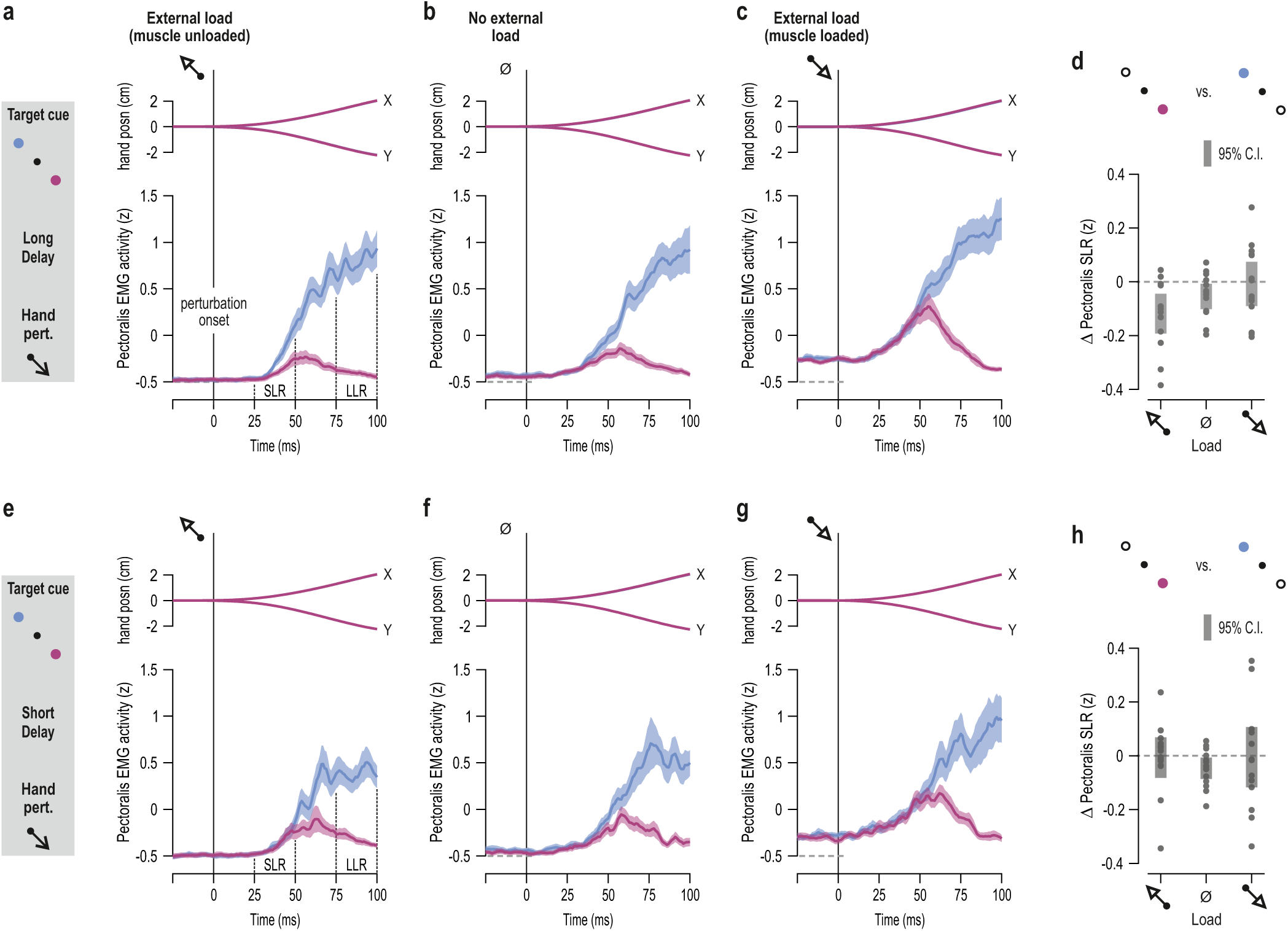
The goal- and delay-dependent modulation of stretch reflex gains is congruent with the preparatory tuning profile of muscle spindles. Mean hand position (posn.) and mean rectified pectoralis EMG activity across participants (N=14) when an external (pre-)load was applied in the direction of pectoralis shortening (**a**), when there was no external load (**b**; but note increased EMG levels prior to time ‘0’ due to co-contraction), and when an external load was applied in the direction of pectoralis stretch (**c**). Shading represents ±1 s.e.m. Data are aligned to the onset of the haptic perturbation (time ‘0’). As the schematic on the far left indicates, the data represent trials where the preparatory delay was relatively long and the subsequent perturbation stretched the pectoralis. SLR denotes the epoch associated with the spinal stretch reflex and LLR the epoch associated with the long-latency stretch reflex or “R3” (for LLR analyses see Results and Fig. 7). **d** Difference in mean pectoralis EMG activity (purple minus blue) in the spinal SLR epoch, corresponding to the data shown in ‘a-c’. Dots represent individual participants and thick vertical lines represent 95% confidence intervals. The SLR of the unloaded pectoralis is suppressed in a goal-dependent manner (‘a’), this relative suppression effect remains but weakens when the muscle is loaded by self-imposed co-contraction (‘b’) and goal-dependent modulation of SLR disappears when the muscle is strongly pre-loaded (‘c’). It is known that SLR responses are affected by load-based or “automatic” gain-scaling but goal-dependent LLR responses are largely not affected by background load. Indeed, here there are consistent effects of goal and delay length on LLR responses regardless of load condition (see also Fig. 7). **e-h** As top row of panels but representing trials where the preparatory delay was relatively short (0.25 sec).

It is already well-established^22-25^ that LLR (or “R3”) responses are goal-dependent and influenced by proprioceptive feedback. Indeed, as can be appreciated by visually inspecting the EMG traces of the pectoralis (Fig. 6) or posterior deltoid (Suppl. Fig. 5), LLR responses were congruent with the goal-dependent afferent results: there is a relative suppression of gains when the target cue is associated with stretch of the particular muscle. But analyses of LLR responses confirmed an even closer connection to the afferent suppression pattern. Specifically, across all load conditions, there was a stronger goal-dependent suppression of LLR responses following a ‘long’ rather than a ‘short’ preparatory delay, with t(13)=-3.63 and p=0.003, t(13)=-3.45 and p=0.004, t(13)=-3.2 and p=0.007, for the pectoralis, anterior and posterior deltoid muscle, respectively (Fig. 7a). Crucially, the ‘short’ preparatory delay here was 250 ms. This is longer than the previously identified delays of 100-150 ms, after which there is apparently no significant improvement in goal-dependent modulation of LLRs, at least as demonstrated by applying perturbations about the elbow joint^26^ alone. In contrast, the current effect of delay length on LLRs is congruent with the spindle tuning profile (i.e., a goal-dependent modulation that is weak early on but markedly stronger late in preparation), hence supporting the presence of an independent and relatively slow-evolving mechanism acting on proprioceptors during reach preparation (as reflected in Fig. 2b-c). Interestingly, our analyses found no significant effect of delay length on LLRs of biceps and triceps muscles (all p>0.05). Performing the same analyses only across cases where the muscles were loaded (i.e., load applied in the direction of muscle stretch) produced equivalent positive findings for shoulder muscles (Fig. 7b), and again no significant effects for ‘elbow’ muscles. However, increasing the workspace of the center-out task in a third experiment (i.e., increasing task demands) reproduced the main positive findings of experiment ‘2’ and also revealed an effect of delay length on LLR responses of elbow muscles.

**Fig. 7:**
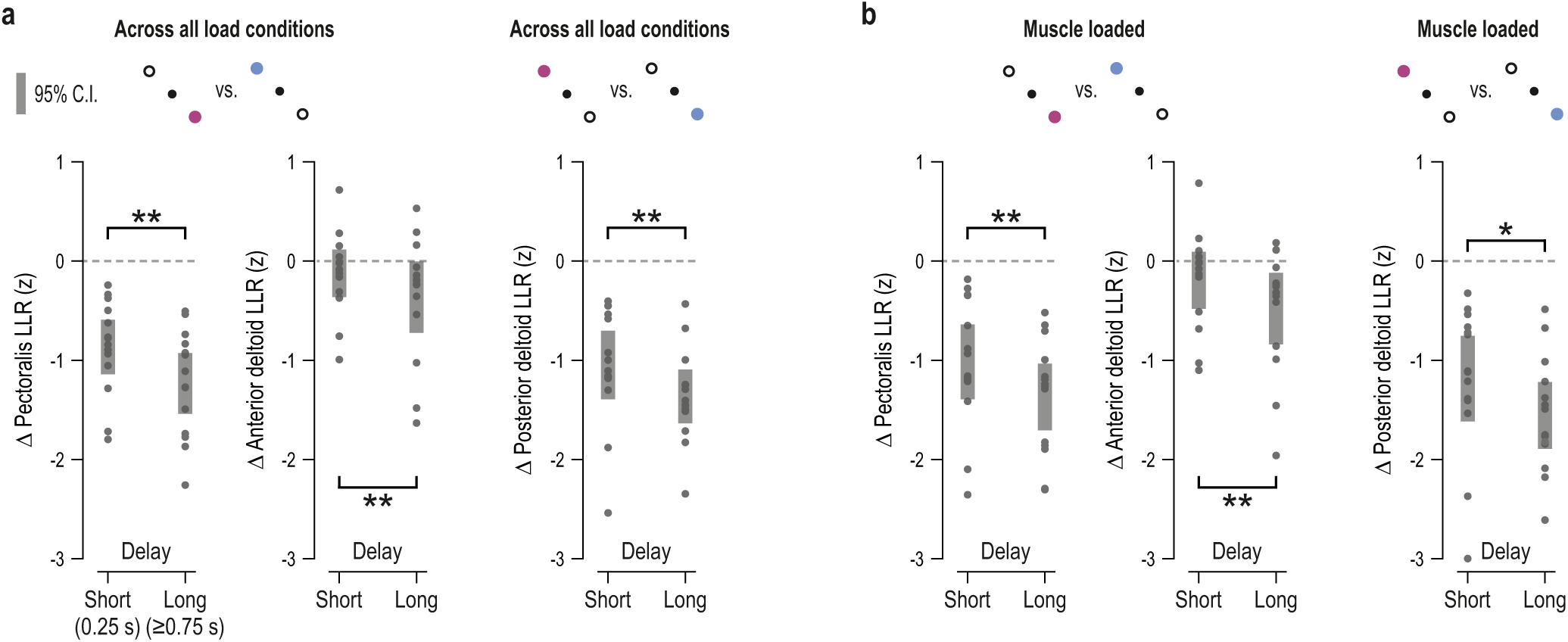
LLR gains reflect the stronger goal-dependent suppression of spindle signals observed at the latter parts of the preparatory delay. Goal-dependent difference in EMG responses of all recorded shoulder muscles at the LLR epoch (e.g., see Fig. 6), with regard to the relatively ‘short’ (250 ms) and ‘long’ (≥750 ms) preparatory delays used in experiment ‘2’. More negative values indicate stronger goal-appropriate behavior (i.e., relative suppression of stretch reflex gains for muscles that must stretch when reaching the cued target). Throughout, each data point represents the average value of a different participant (N=14) and thick vertical lines represent 95% confidence intervals. Stars indicate p values following a within-measures t-test, with double stars indicating p<0.01 and single p<0.05. These results demonstrate a weaker goal-dependent modulation of LLRs when the preparatory delay is ‘short’, regardless if contrasted across all load conditions (**a**), or only for the cases where the muscle was externally loaded i.e., the load was applied in the direction of muscle stretch (**b**). The ‘short’ delay here was 250 ms, which is substantially longer than the previously reported minimum delay for inducing full expression of goal-dependent LLR responses following perturbations about the elbow (i.e., 150 ms; see main text). In contrast, the effect of delay length on LLR gains is congruent with the temporal evolution of spindle suppression (i.e., significantly stronger at the latter parts of preparation; Fig. 2c).

Specifically, a third experiment implicating a larger number of visual targets produced equivalent results for pectoralis SLR (i.e., experiment ‘3’; Fig. 8). When the preparation delay was relatively long and the pre-load was in the direction of pectoralis shortening (Fig. 8a-c), there was a goal-dependent suppression of SLR gains, with t(11)=-4.1, p=0.002 (Fig. 8d). Although for most participants SLRs were suppressed in the no-load condition as well (middle column in Fig. 8d), the overall difference was deemed not significant (p=0.2; note the one deviant value >0). As in experiment ‘2,’ there was a small but significant suppression of SLRs in the no-load condition when the delay was short (Fig. 8e-h), with t(11)=-2.8, p=0.017. As in experiment ‘2’, LLRs of shoulder muscles closely reflected the spindle suppression pattern. That is, across all load conditions, goal-appropriate suppression of shoulder muscle LLR was stronger if a ‘long’ rather ‘short’ delay preceded congruent perturbations, with t(11)=-2.42 and p=0.034, t(11)=-2.22 and p=0.048, t(11)=-2.3 and p=0.042, for the pectoralis, anterior and posterior deltoid muscle, respectively. A significant effect of delay length was also found for the triceps lateralis, with t(11)=-3.74 and p=0.003. To better contrast the seemingly conflicting results of experiment ‘2’ and ‘3’ regarding the effect of preparatory delay length on ‘elbow’ muscles, the data were contrasted separately for each of the three main axes of motion involved in experiment ‘3’: diagonal (as in experiment ‘2’), vertical and horizontal (Fig. 9). As in experiment ‘2’, there was no effect of delay length on biceps and triceps LLR responses when action was required along the diagonal axis (p>0.05). However, along the horizontal dimension, there was a stronger goal-dependent suppression of biceps LLR following a long delay (t(11)=-2.73 and p=0.02), and the same effect was evident for the triceps when target cues required action along the vertical axis (t(11)=-4.02 and p=0.002). The above results suggest that the increased task demands (i.e., larger workspace) of experiment ‘3’ necessitated the proprioceptive control of a larger group of muscles, even though the perturbations themselves where always applied along the diagonal axis, as in experiment ‘2’. Taken together, the results of experiment ‘2’ and ‘3’ confirm that the spindle tuning profile observed during preparation (i.e., a goal-dependent modulation that was weak early on but markedly stronger late in preparation) impacted stretch reflex gains, particularly those of LLRs that are largely unaffected by “automatic” gain-scaling”^19-21^ and generally can contribute much more to generated forces than SLRs.

**Fig. 8:**
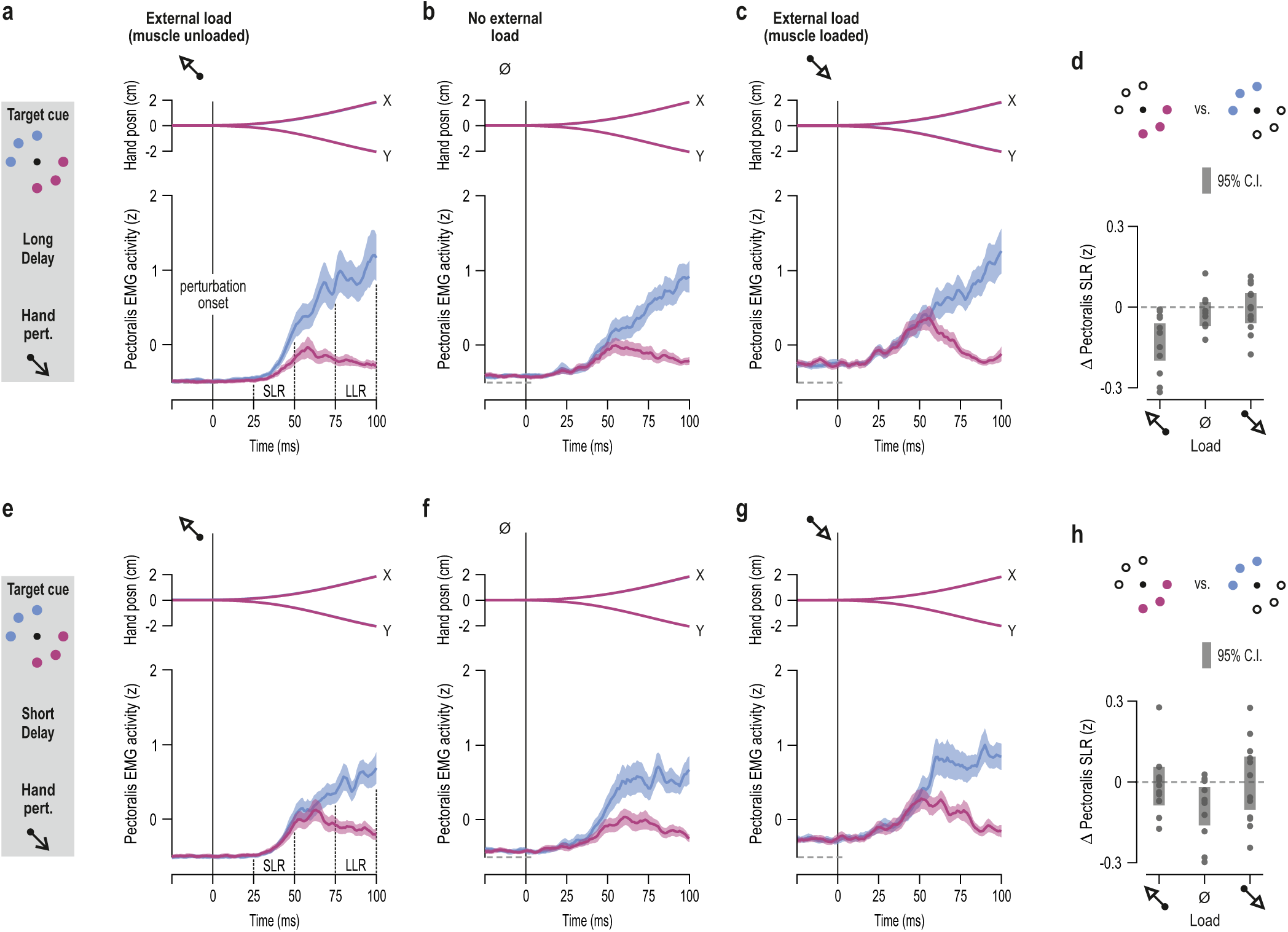
Third experiment utilizing a larger workspace also demonstrates congruent preparatory tuning of stretch reflex gains. Experiment ‘3’ was conducted as per experiment ‘2’ (Figs. 5-6) except in this case six targets were employed rather than two (see left schematics) and the long and short preparatory delays were 1.2 and 0.2 sec, respectively. **a-c** Mean hand position (posn.) and mean rectified pectoralis EMG activity across participants (N=12), when an external load was applied in the direction of pectoralis shortening (**a**), when there was no external load (**b**; but note increased EMG levels prior to time ‘0’ due to co-contraction), and when the load was applied in the direction of pectoralis stretch (**c**).The data represent trials where the preparatory delay was relatively long and the subsequent perturbation stretched the pectoralis. Shading represents ±1 s.e.m. SLR denotes the epoch associated with the spinal stretch reflex and LLR the epoch associated with the long-latency stretch reflex (see main text for LLR analyses). **d** Difference in mean pectoralis EMG activity (purple minus blue) in the SLR epoch, corresponding to the data shown in ‘a-c’. Dots represent individual participants and thick vertical lines represent 95% confidence intervals. **e-h** As the top row of panels but representing trials where the preparatory delay was relatively short.

**Fig. 9:**
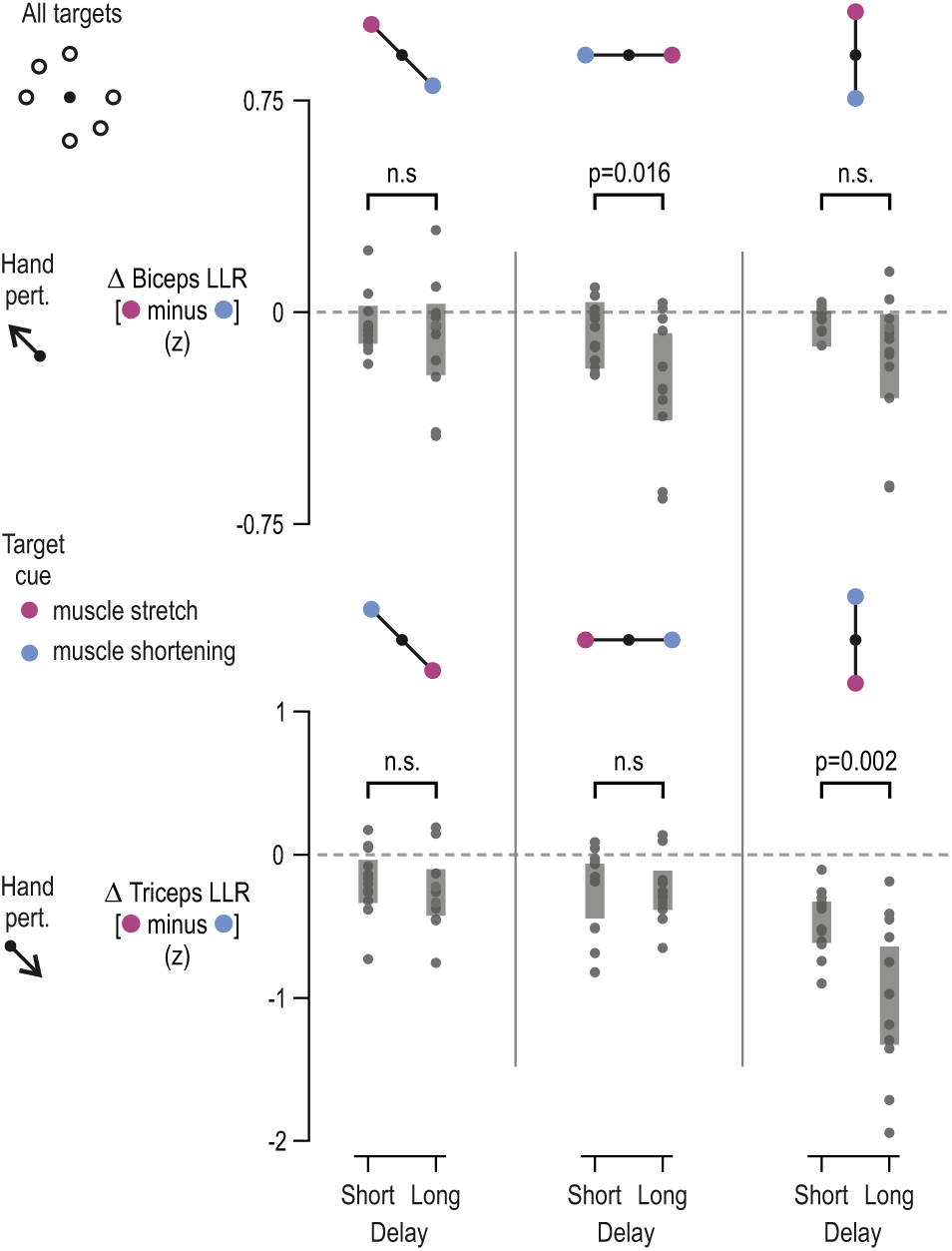
LLR gains of biceps and triceps are also suppressed as a function delay when a larger workspace is involved. As Figure 7, but here the z-normalized EMG data originate from Experiment ‘3’, where six targets were used (i.e., three axes of motion: vertical, horizontal and diagonal). The data are collapsed across all load conditions. More negative values indicate stronger goal-appropriate behavior (i.e., relative suppression of stretch reflex gains for muscles that must stretch when reaching the target). Throughout, each data-point represents a different participant and thick vertical lines represent 95% confidence intervals. p values resulted from within-measures t-tests. As the case in ‘Experiment ‘2’, the LLR responses of biceps and triceps were not significantly different as a function of delay length when preparing to reach targets along the diagonal axis (left column; only axis used in Experiment ‘2’). However, such effects are observed for the biceps brachii and triceps lateralis when preparing to act along the horizontal axis (middle column) and vertical axis (right column), respectively. This suggests that the larger workspace employed in experiment ‘3’ (vs. ‘2’) induced goal-dependent proprioceptive control of a larger group of muscles, including elbow flexor and extensor muscles. The EMG traces pertaining to the significant triceps result are shown in Supplementary Figure 6.

## Discussion

Our results indicate that movement preparation can involve goal-appropriate tuning of muscle spindle receptors. This assigns a novel and specific role to central preparatory activity in proprioceptive tuning with implications for stretch ‘reflex’ behavior. Our findings are congruent with classic results concerning preparatory activity in the CNS and its two hallmarks which are (a) that preparatory activity should not overtly affect concurrent skeletal muscle activity and, (b) preparatory activity needs to somehow promote or facilitate the planned movement. The current study helps bridge the gap between traditional views where preparatory activity is seen as representing specific movement parameters^1-4^, and the more recent claims that movement preparation shapes an initial state of a dynamical system whose evolution produces the planned movement^5,6^. We show that such an ‘initial’ state may partly pertain to the state of the peripheral proprioceptive apparatus, which can predispose the system for sensory attenuation and stretch reflex suppression of muscles that will lengthen during the desired voluntary movement. One can speculate that failure to properly engage this mechanism may contribute to target undershoot (‘dysmetria’) and perhaps spasticity. But even in the healthy population, this preparatory mechanism may represent a significant source of individual differences in motor performance. Indeed, we found that higher levels of tonic type Ia discharge at late preparation are associated with worse reaching performance (i.e., larger delays in attaining peak velocity; Fig. 4b). This relationship can be understood in terms of the spindle’s role in generating negative feedback via stretch reflexes.

The current study is the first to record muscle afferent responses during movement preparation (i.e., over a dedicated delay period) in a context where voluntary reaching movements were actually made. One other study^27^ implicating the upper limb looked at spindle responses when anticipating the need to make a contraction that would oppose an expected external perturbation. No preparatory effects were found but we believe our paradigm better reflects the state of affairs when reaching in every-day life, as the task combined true reaching intention and action. There has also been strong evidence of preparatory activity in spinal interneurons^28,29^, but our study is the first to document preparatory changes in sensory elements of the peripheral nervous system. Hence, we show that preparatory activity can be associated with implementation of control policy. The central origin of fusimotor control was not directly addressed by the current study, but some clues are offered. Preparatory activity in the cortex can appear in as little as 50 ms following onset of the target cue^6,30^ and activity of corticospinal neurons can be suppressed during movement preparation^31^. Although the spindle suppression effect seems to begin early, at ∼80 msec after the target cue, the suppression is initially weak and becomes much stronger at the later stages of preparation (Fig. 2b-c). The evolution of this effect is congruent with the tendency of preparatory CNS signals to differentiate more clearly according to preferred direction closer to the onset of the ‘Go’ cue e.g.,^5,6^. The general profile of proprioceptive suppression suggests an early and a later phase to this process (Fig. 2b-c). This may in turn reflect two different sources of fusimotor control, an early subcortical one (e.g., brainstem^32-34^) and a later cortical one. However, identifying the precise CNS origin or specific descending pathways associated with preparatory fusimotor control is unnecessary for the purposes and main novel claim of this study i.e., that there is advantageous preparatory tuning of muscle spindle receptors.

The general expectation of no specific role for spindle receptors during movement preparation has been formulated indirectly, primarily through behavioral studies examining spinal SLR responses in surface EMG from the upper limb. Although there has been some evidence of goal-dependent modulation of SLRs, both at the level of digits^35^ and at more proximal areas^36^, previous studies at the level of the digits and more recent studies using robotic platforms to assess reflex responses of more proximal muscles have not identified goal-dependent spinal SLR responses; but such responses are consistently found at transcortical latencies^22^. The results of experiments ‘2’ and ‘3’ suggest that, given a particular experimental design, goal-dependent modulation of SLRs can be consistently induced (Figs. 6, 8 & Supplementary Fig. 5). Two important elements of our experimental design are the systematic manipulation of background load and ensuring that movement control is required for reaching every target. Regarding the first, many previous studies either did not account for the background activation levels of muscles or deliberately pre-loaded muscles to ensure detectable levels of surface EMG in the SLR epoch. We show that strongly loading a muscle can potentially obscure evidence of goal-dependent proprioceptive tuning (e.g., Fig. 6a-c). That is, our results show that load-related or “automatic” gain-scaling^19-21^ of SLRs for the purposes of postural control may compete or otherwise interfere with target-dependent tuning of spinal SLRs. But muscle spindle gains-modified by independent γ control-are not necessarily affected by background mechanical loading. Indeed, when imposing stretch of the isometrically loaded radial wrist extensor, no clear net difference in spindle sensitivity is found, as an approximately equal number of ‘dynamic’ and ‘static’ fusimotor effects appear with these two having opposite effects on spindle gain^37^. Nevertheless, across all load conditions, we show a modulation LLR gains that closely reflects the spindle’s preparatory tuning profile (e.g., Fig. 7). That is, we show a goal-appropriate modulation of gains that is stronger following longer than relatively shorter preparatory delays.

Crucially, the ‘short’ delays used in our study are longer than the previously identified delays of 100-150 ms, after which no significant improvement in goal-dependent modulation of LLRs has been previously reported, at least as demonstrated by applying single-joint perturbations about the elbow^26^ (in fact, they noted a deterioration of LLR behavior for delays >200 ms^26^). The LLR results stemming from the center-out reaching paradigm of the current study are supportive of an independent and relatively slow-evolving mechanism acting on proprioceptors during reach preparation. Our findings are also compatible with reported improvements in reach movement quality that occur for preparation delays >150 ms^10^. Equivalent SLR effects were also detected in the current study, except when muscles were strongly pre-activated. A likely reason for the apparent saturation of spinal SLR responses in loaded muscles is the excitation state of spinal circuits. Even if spindle gains remain suppressed (but not fully) when about to stretch a loaded muscle, a high spinal excitability level can still lead to large SLR responses, obscuring any goal-dependent tuning of muscle spindles (i.e., ceiling effect; but the opposite extreme is also problematic, see e.g., Supplementary Fig. 5c). Nevertheless, it is well-known that LLR responses are robustly goal-dependent and largely immune to load-based or “automatic” gain-scaling”^19-21^. In this context, appropriate LLR responses can emerge simply by linking goal-dependent afferent signals to transcortical feedback circuits^22,38,39^ that are not subject to automatic gain scaling.

One study that examined stretch reflex responses of unloaded (“pre-inhibited”) elbow muscles following perturbations of the forearm about the elbow did not report task-dependency of SLR, instead documenting flat-lined EMGs at spinal latencies^40^. But such null SLR responses across targets could have possibly been due to high levels of reciprocal inhibition, brought on by placing a large ‘pre-inhibiting’ load at the single joint; in the congruent perturbation case in particular, “no substantial muscle activity” was observed even at voluntary latencies. Indeed, their ‘IN-OUT’ paradigm did not require participants to engage in movement control during congruent perturbations as the hand was moved inside the target area early-on by the perturbation itself (i.e., large target area was adjacent to hand origin). A second important element of our experimental design was ensuring that active/voluntary movement control was required for reaching the target throughout the task. That is, participants had to actively complete movement to the target on every trial, including after congruent perturbations (see e.g., purple velocity profiles in Fig 5c-e). Hence, on such trials the participants were implicitly encouraged to ‘facilitate the reach’ rather than ignore or ‘not resist’ a perturbation. The intent to actively engage in attaining a movement’s goal may be necessary for invoking preparatory proprioceptive control. Overall, our multi-joint reaching paradigm reflects more closely the classic center-out delayed-reach paradigm where targets are placed at some distance from the hand. Very recently, it has been shown that elbow-actuating muscles exhibit SLR responses that are tuned to the position of the hand relative to a single target, rather than the state of the muscles themselves^41^. Although this does indicate a higher level of sophistication by spinal monosynaptic circuits than previously thought, the differing SLR responses were a function of the different configurations of the limb, whereas the goal of the task (i.e., location of the target) remained the same across experimental conditions. In contrast, in our study we were able to isolate an effect of external goal (visual target) on stretch reflex gains by varying target location while maintaining the state of the limb constant across conditions of interest.

The current findings highlight that muscle spindle receptors and their independent motor system can serve more decisive and task-dependent roles in sensorimotor control than generally thought. Traditionally, the spindle organ has been seen as a peripheral mechanoreceptor that provides reliable information about a muscle’s kinematic state. An interesting recent proposition is that the mechanoreceptive part of spindles responds best to force-related rather than length-related variables, as shown in passive (‘electrically quiescent’) muscles^15^. Indeed, when performing continuous active sinusoidal movements with a single digit in the presence of external loads, we have also shown that spindle afferent activity from digit extensors best encodes a combination of velocity and net external mechanical force^42^. But our more recent work examining spindle responses in visuomotor learning (i.e., visuomotor rotation) revealed fundamental changes in spindle output as a function of task stage (e.g., encoding position vs. velocity in the ‘washout’ stage), with no fundamental differences in mechanical state across the task’s stages^16^. Besides indicating that the fusimotor system is a specific contributor in visuomotor learning, the aforementioned study showed that spindle output can be modified based on changes in the visual environment alone. This is in line with the findings of the current study (e.g., Fig. 2b-c). Very recent spindle afferent recordings during passive movement of the foot also indicate that visual feedback can affect spindle output^43^. Accumulating evidence therefore suggests that human spindles can transcend their traditionally-ascribed role as mechanoreceptors invariably encoding some muscle state regardless of context or goal. The traditional account also essentially assumes the purpose of fusimotor control is to ensure that spindles keep functioning as a reliable mechanoreceptors, as described by the textbook version of ‘alpha-gamma co-activation’^44^. In cats, it has been shown that spindles can receive a different ‘fusimotor-set’^45^ depending on the behavior the animal is engaged in (i.e., primarily variations of standing or gait in different contexts), but the specific benefit of the different fusimotor sets has been unclear, and these sets generally seem to reflect the alertness state of the animal. Here, we show spindle gain modulation as a function of visually-determined goals within the same behavior (reaching), including evidence of how this spindle tuning can promote motor performance (i.e., Figs. 4-9).

The traditional view of spindles as mundane proprioceptive sensors is the one currently adopted by prevalent computational frameworks of sensorimotor control^46-50^. Part of these suggest that our brain predicts the sensory consequences of action and then compares internal predictions and actual incoming sensory signals (‘sensory cancellation’), with no discrepancy between the two indicating agency of action. With regard to primary muscle spindles in the context of planned reaching movements, our results suggest that the nervous system does more than these computational frameworks describe. Presumably still based on internal models and predictions of future outcomes given an intention or goal^49,50^, the system seems able to proactively choose and implement a change in sensory feedback gains at source (e.g., Fig. 2-3). That is, in planned voluntary reach, the ‘controller’ can proactively modify the ‘plant’ (i.e., adjust sensitivity of the plant’s sensors) in order to prevent consequences (negative feedback) that would otherwise interfere with execution of the intended action. Beyond its role in planned reaching, the independent and direct control of sensors via γ motor neurons may well constitute an important overarching third dimension in sensorimotor control, in addition to (i) top-down processes leading to α motor neuron control and, (ii) the selective gating and internal processing of sensory signals. Understanding the full potential and implications of this neglected third dimension in active behaviors will be a major focus of our future work. By demonstrating advantageous tuning of spindles in movement preparation, the current study supports the notion of a ‘third way’ in which the nervous system can exert goal-dependent sensorimotor control.

## Methods

### Human participants

We recorded afferent activity from 9 adults in the first experiment (mean age of 27 and SD = 3 years; 5 were male), 14 individuals took part in the second experiment utilizing a robotic manipulandum platform (mean age of 24.5 and SD = 4 years; 6 were male), and an additional 12 adults participated in the third experiment utilizing the same platform (mean age = 25 and SD = 5 years; 5 were female). All participants reported having no motor or cognitive disabilities, had normal or corrected vision, gave their written consent before taking part and were financially compensated. The current experiments were part of research programs approved by the Regional Ethics Committee of Umeå and followed the Declaration of Helsinki regarding research with humans.

### Experimental setups

#### Microneurography platform

The participants were seated reclined on an adjustable chair with their right forearm resting on a cushion. The activity in single afferents from wrist or digit actuator muscles was recorded along with wrist joint kinematics and EMG activity from relevant forearm muscles (Fig. 1a). Participants used their right hand in order to perform a classic center-out reaching task, where each target is first cued before a ‘Go’ cue to move is issued (the task is described in more detail below). A clamp proximal to the wrist stabilized the upper arm and helped prevent electrode dislocations, but hand movements about the wrist were fully unrestrained in this setup. In ‘classic’ center-out reaching tasks, target location is normally presented on a monitor and so is the visual feedback on the location of the hand, represented by a moving cursor. The approach was the same here: visual feedback was provided by a monitor that was placed across from the participants and elevated at about their eye-level. They controlled the 2D location of a cursor on the monitor through wrist movements recorded by a FASTRAK® sensor attached to the dorsal surface of the hand with double-sided tape. The initial posture of the hand represented a neutral wrist position which in turn corresponded to the ‘origin’ position of the cursor (Fig. 1a). In this neutral position, the hand (e.g., third metacarpal joint) was aligned with the long axis of the forearm, and to hold this position against gravity the participants had to produce a constant low-level contraction mainly in the extensor carpi radialis. Wrist radial/ulnar rotations controlled cursor movements in the vertical visual axis and flexion/extension controlled cursor movements in the horizontal axis (Fig. 1a). One degree movement at the wrist corresponded to 0.7 cm on-screen movement of the visual cursor. Visual targets not involved in an ongoing trial where represented as light brown circle outlines (1.5 cm radius; origin outline had 1 cm radius). The targets were placed symmetrically around the origin in 45° intervals so that movements in all major directions were induced (Fig. 1a). The distance between the center of the origin and the center of a target was 12°, but a minimum wrist movement of 10° was required for successfully reaching from origin to target (i.e., edge to edge).

#### Robotic platform

Here the participants were seated upright on an adjustable chair and their right hand grasped the handle of a robotic manipulandum (Fig. 5a; KINARM end-point robot, BKIN Technologies, CA). Although not displayed in Figure 5a, the participant’s right forearm was placed inside a thin cushioning foam structure attached to a custom-made airsled; this structure supported the participant’s forearm and allowed frictionless movement of the arm in a 2D plane. A piece of leather fabric with Velcro attachments was wrapped tightly around the forearm and hand, reinforcing the mechanical connection between the airsled, the handle and the hand. This attachment also fixated the hand so it remained immobile about the wrist and straight (i.e., aligned with the forearm) throughout the experiment. The forces exerted by the participant’s right hand were measured by a six-axis force transducer (Mini40-R, ATI Industrial Automation) embedded in the handle, and the system also generated kinematic data with regard to the position of the handle. The KINARM also produced controlled forces on the hand, both for the background (pre-) loading of muscles and for creating position-controlled mechanical perturbations. Surface EMG was concurrently recorded from seven muscles actuating the right arm (see the relevant section for more details). Visual feedback was very similar to that presented in the microneurography experiment, but in the robotic platform visual stimuli were displayed in the plane of movement by way of a one-way mirror, on which the contents of a monitor were projected. The participants had no direct vision of their hand (Fig. 5a), but position of the hand was visually represented by a white dot (‘cursor’; 1 cm diameter). Targets not involved in an ongoing trial where displayed as circle outlines (1.2 cm radius; origin outline had 0.65 cm radius). The targets were placed symmetrically at a distance of 9 cm from the origin.

### Specific experimental procedures

#### Microneurography – hand movement task

In the behavioral task associated with microneurography, the participants (n=9) were instructed to place the cursor inside the origin circle and wait there immobile before a trial could start. After a random wait period (0.5 - 2.5 sec), one of the eight different targets would suddenly turn from a circle outline to a filled red circle of the same size. This indicated which target the participant had to reach once the ‘Go’ cue appeared. The ‘Go’ cue in this case was the red target suddenly turning into a green outline of the same size. For the majority of the participants (7/9), the time between onset of the target cue (red circle) and onset of the ‘Go’ cue was a fixed 1.5 sec (‘preparatory period’). To assess whether any major afferent firing patterns during movement preparation were critically sensitive to major characteristics of the particular preparatory period, we used 1 sec as the preparatory period with one participant, and 1.5 sec + random time (1-500 ms) for another. No substantial differences in firing patterns were found between these afferents and the rest Ia. To aid subsequent analyses, data from the initial 1.5 sec were used in the latter case, and in the former case the data during the 1 sec periods were resampled offline to 1.5 sec. In all experiments, the participants were instructed to initiate the reach movement promptly upon onset of the ‘Go’ cue and to move at a naturalistic speed. To promote this behavior, participants received visual feedback on their performance upon reaching a target. That is, they received the message “Good” if they managed to reach the target within 1 sec following onset of the ‘Go’ cue and “Fail” if they took longer. After receiving feedback, the participants returned to the origin to initiate the next trial. The task continued until the afferent recording was lost due to an accidental dislocation of the electrode, an all too common occurrence when recording during naturalistically fast active movement (but at least 24 trials i.e., three blocks of trials were recorded with each afferent; see below for more details). Trials where movement was initiated prematurely (i.e., before the ‘Go’ cue) were excluded from analyses, but these represented just one trial per afferent on average, and in no case more than two trials per recorded afferent. To familiarize the participants with the center-out task and promote good performance at it during microneurography, they practiced the task for ∼10 minutes before microneurography began.

#### Robotic platform – arm movement tasks

Two experiments where conducted using a robotic platform (experiment ‘2’ and ‘3’), with each experiment employing a different set of participants. Before the main task of either experiment, each participant initially performed a brief unperturbed center-out reaching task that was very similar to that during microneurography. This introductory task was included in order to establish a closer link between the behavioral task in microneurography and the main task applied with the robotic platform (described below). Specifically, in this brief center-out task, participants were instructed to bring the hand in the origin circle and remain there immobile. After a wait period of one sec + random time (1-500 ms), one of the eight peripheral targets/outlines turned into a filled red circle of the same size, indicating which target the participant had to reach once the ‘Go’ cue appeared (‘Go’ = target turning green). The ‘preparatory period’ here was a fixed 1.5 sec, to match the case during microneurography. Participants had to move at a naturalistic speed and upon reaching a target they received visual feedback on their performance. Counting from the onset of the ‘Go’ cue, the feedback was “Too Slow” if the reach movement lasted >1400 ms, “Too Fast” if <400 ms, and “Correct” if the movement duration was in-between the two stated extremes. After receiving feedback, the participants returned to the origin to initiate the next trial. There were 80 trials in total (i.e., 10 repetitions x 8 targets), presented in a block-randomized manner, with one set of eight different targets representing a ‘block’. The task lasted ∼5 minutes.

Following a short break of a few minutes, the participants then performed the main behavioral task. In experiment ‘2’ (e.g., Figs. 5-6), the main task lasted for ∼1 hour, whereas in experiment ‘3’ (e.g., Fig. 8) the task lasted ∼1.5 hours. The main task was designed to emphasize reflex responses from shoulder actuators, allowing the possibility to extend positive findings to the most proximal areas of the upper limb, although elbow muscle reflexes where also stimulated. Specifically, visual feedback in the main behavioral task of experiment ‘2’ was the same as in the brief introductory task described above, except that two rather than eight targets were employed (Fig. 5a-b) and the cursor position was frozen for the duration of haptic perturbations. Before each trial begun, the participants brought the hand (i.e., cursor) inside the origin circle. After a wait of one sec + random time (1-500 ms), the robotic arm was programmed to elicit a slow-rising 4N load (rise-time 800 ms, 1200 ms hold-time) in the front-and-left direction (‘135°’ direction) or right-and-back direction (‘315°’ direction), or no load was applied. A substantial load could therefore be present at this point in each trial, with the function of pushing towards one or the other target (Fig. 5a-b). Because the participants were instructed to maintain their hand in the middle of the origin circle during this phase of the trial, the ultimate purpose of this maneuver was loading/unloading of the recorded actuators, primarily the posterior deltoid or pectoralis and anterior deltoid. After an additional 1.2 sec where the full force of the load was countered while the hand remained still, one of the targets was cued by becoming a red filled circle. After a preparatory period of either 0.25, 0.75 or 1.25 sec, a position-controlled perturbation of the hand occurred (3.5 cm displacement, 150 ms rise time, no hold period), swiftly moving the hand towards the middle of one or the other group of targets i.e., in the 135° or 315° direction. The specific preparatory delays were chosen to match the middle of epochs ‘1-3’, as identified in Fig. 2b-c. The haptic perturbations were designed to induce the kinematics of a fast naturalistic point-to-point movement (i.e., approximate bell-shaped velocity profile; e.g., see Fig. 5c-e) and promote stretch reflex responses primarily in shoulder muscles. The robot was allowed to employ maximum available stiffness (∼40,000 N/m) -if necessary- to achieve the desired kinematics on every trial. The KINARM robot was able to reliably impose the required hand kinematics during these perturbations regardless of background load/force conditions. When the haptic perturbation ended (i.e., 150 ms after perturbation onset), the ‘Go’ cue suddenly appeared and the participants swiftly reached to this highlighted target. The trial ended when the participants kept their hand immobile inside the target for 0.3 sec, after which they received visual feedback on their performance (i.e., “Correct”, “Too Fast” or “Too Slow”), as per the brief introductory task. The participants then returned their hand to the origin to initiate the next trial. Each block of trials represented one repetition of each level of each condition (i.e., block = 36 trials: 2 targets x 2 perturbation directions x 3 preparatory periods x 3 load conditions) and there were 15 repetitions of the complete trial block; that is, the total number of trials was 540. The trials were presented in a block-randomized manner, and therefore all perturbations were unpredictable to the participants in terms of their timing (onset) and direction. The participants had the opportunity to take a short break at the end of each block of trials. ‘Experiment 3’ was essentially the same as ‘Experiment 2’ except that six targets were used rather than two, and the two preparatory delays were 0.2 and 1.2 sec, also referred to as ‘short’ and ‘long’. Each block of trials represented one repetition of each level of each condition (i.e., block = 72 trials: 6 targets x 2 perturbation directions x 2 preparatory periods x 3 load conditions) and there were 10 repetitions of the complete trial block; that is, the total number of trials was 720.

### Muscle afferent recordings

Single spikes in afferents originating from either the radial wrist extensor (*extensor carpi radialis*), the ulna wrist extensor (*extensor carpi ulnaris*) or the common digit extensor (*extensor digitorum communis*) were obtained using the technique of microneurography^51^. The radial nerve of the right arm was targeted, and isolated single action potentials were categorized as originating from spindle or Golgi tendon organ afferents following standard procedures described in detail elsewhere^17,18,42^. In total, 12 muscle spindle afferents (8 ‘type Ia’ and 4 ‘type II’) and 3 Golgi tendon organ afferents were recorded from 9 participants (minimum of one recorded afferent per included participant). With all afferents a minimum of 24 movement trials were recorded (i.e., 3 repetitions of a movement direction) and with some the recording lasted longer, allowing for more repetitions to be sampled.

As expected, the primary spindle afferents responded with higher overall firing rates to dynamic muscle stretch than muscle shortening. Just one afferent from a digit actuator was not responsive to one of the three ‘stretch’ target directions (i.e., upper left direction) but was very responsive to the other two stretch directions. Likely causes for such variability include the particular set of fusimotor supply and the precise location of the spindle organ inside the muscle. The number of afferents recorded in this study reflects that in previous studies examining single afferent activity during active movement e.g.,^16,42,52^. Moreover, it has been shown that a small number of spindle afferents can provide a reliable representation of the firing patterns observed in the underlying afferent population e.g.,^14^. This is not surprising, as all muscle spindle organs are placed mechanically “in parallel” with the skeletal muscle fibers, and the spindle acts as an integrator of activity from multiple fusimotor fibers.

### Muscle EMG recordings

In the microneurography experiment, custom-build surface electrodes (Ø 2 mm; 12 mm apart) were used for recording EMG from the common digit extensor and digit flexor muscles, as well as from the four main wrist actuators (extensor carpi radialis, extensor carpi ulnaris, flexor carpi radialis and flexor carpi ulnaris). The location of each electrode on the forearm was chosen using a hand-held stimulator probe and isometric contraction/relaxation maneuvers. In experiment ‘2’ and ‘3’, the Delsys Bagnoli system (DE-2.1– Single Differential Electrodes) was used to record surface EMG from the pectoralis, posterior deltoid and the anterior deltoid. We also recorded EMG from the brachioradialis, biceps and triceps areas. In all experiments, EMG electrodes were coated with conducive gel and attached to the skin using double-sided tape.

### Data sampling and processing

The data generated during the microneurography experiment were sampled digitally using SC/ZOOM^™^. Single action potentials were identified semi-automatically under visual control. The EMG channels recorded during microneurography were root-mean-square processed with a rise-time constant of 1.0 ms and a decay-time constant of 3.0 ms; they were then digitally sampled at 1600 Hz. The EMG channels were high-pass filtered with a fifth-order, zero-phase-lag Butterworth filter with a 30 Hz cutoff. Kinematic and force data from the KINARM platform were sampled at 1 KHz. The recorded EMG signals were band-pass filtered online through the Delsys EMG system (20-450Hz) and sampled at 2 kHz. This EMG data was also high-pass filtered with a fifth-order, zero phase-lag Butterworth filter with a 30 Hz cutoff and then rectified. To be able to compare and combine EMG and afferent data across muscles and participants, the raw data were normalized (z-transformed), similar to the procedure described elsewhere^16,36,42,53^. Briefly, for each individual muscle (or individual afferent), all relevant raw data traces were concatenated, and a grand mean and standard deviation was generated. These two numbers were then used to produce the normalized ‘raw’ EMG data for each muscle or produce the normalized firing rate of each afferent (i.e., by subtracting the grand mean and then dividing by the standard deviation). Exemplary untreated raw data are also presented (Fig. 1b-c). For plotting purposes alone, continuous firing rate signals were smoothed using 10 ms moving window (i.e., Fig. 2b) and a 5 ms moving window was used for EMG signals (e.g., Fig. 6). Throughout, data tabulations were performed using Matlab® (MathWorks, Natick, MA, USA).

### Procedures for statistical analyses

The main statistical approach involved conducting repeated-measures t-tests and ANOVA, and complementary planned comparisons on kinematic, EMG and normalized spindle firing rate data observed during the preparatory periods (experiment ‘1’), and single sample t-tests on EMG data pertaining to spinal and long-latency stretch reflex responses elicited during haptic perturbations (experiment ‘2’ & ‘3’). Specifically, with regard to the analysis of the afferent data, it is known that kinematic variables such as position (i.e., muscle length) and its derivatives as well as spindle-bearing EMG activity can affect spindle output, with muscle velocity (i.e., first derivative of muscle length) believed to normally exert the largest influence. To generate estimates of muscle length (tendon excursion) from the recorded wrist angular data we used established physiological models^54,55^ as done previously elsewhere^16-18^. The impact of kinematic and EMG variables on primary spindle afferent output during movement was examined by performing a forward step-wise regression using population signals (i.e., grand mean of median responses from single participants/neurons; Fig. 3). However, as expected, kinematic and EMG variables represented very small levels of variability during the main period of interest (i.e., immobile hand during the preparatory period; Supplementary Fig. 1). The main analyses of data from ‘Experiment 1’ examined potential effects of the goal/target of each trial (i.e., prospective movement direction: muscle stretch vs. shortening) during movement preparation, and no systematic variation in kinematic variables or EMG was found as a function of goal (Supplementary Fig. 2).

To investigate the impact of goal we grouped different trials into those associated with clear stretch vs. clear shortening of the spindle-bearing muscle (Fig. 2a) based on the aforementioned physiological models, but this grouping is nevertheless intuitive and straightforward (e.g., for the radial wrist extensor, targets requiring wrist flexion and/or ulna deviation were classified as ‘muscle stretch’ targets). For each single afferent, the normalized raw data across trials were first aligned to the onset of the target cue. To more clearly isolate possible changes in firing rate as a function of target, the median firing rate observed during the 0.5 sec period before target onset (‘baseline’) was subtracted from the entire firing rate signal on a trial by trial basis. The firing rate signals were collapsed across trials in order to get a single averaged (median) response signal for each afferent and target group (i.e., ‘stretch’ vs. muscle ‘shortening’ targets). Averaging across all afferent signals for each target group gave an estimate of population responses (Fig. 2b). From each averaged afferent signal, the data points used in statistical tests (ANOVA / t-test) were the median value across each of three epochs of equal length, termed ‘Epoch 1’, ‘2’ and ‘3’ (Fig. 2b-c). The data-points pertaining to individual spindle afferents (i.e., Fig. 2c) were entered into a two-way repeated-measures ANOVA, of the design 2 (goal/direction) x 3 (Epoch). Single-sample t-tests, planned comparisons and simple linear correlations were also performed. The same single-sample t-test analyses were also performed with kinematic and EMG data, as described in the Results section.

With regard to stretch reflex responses to haptic perturbations (i.e., experiment ‘2’ & ‘3’), the analyses focused on established time-periods known to reflect the output of spinal and supraspinal stretch reflex circuits. Specifically, across all experiments, the onset of movement or kinematic perturbation was defined as the point where movement velocity (i.e., 1^st^ derivative of Euclidean displacement) exceeded 5% of peak velocity during the perturbation phase (note the position-controlled perturbations had an approximate bell-shaped velocity profile). Using the onset of the kinematic perturbation to signify time zero, the spinal stretch reflex response (SLR) is defined as that occurring in the epoch 25 – 50 ms post perturbation, whereas the long-latency reflex response (LLR a.k.a. “R3”) is defined as that occurring in the epoch 75 – 100 ms post perturbation e.g.,^23,41^. The magnitude of the SLR and LLR response was representative of changes in gain, as the same input (perturbation) was provided when the hand was at a common start position. An epoch of the same length as the SR one was used for representing pre-perturbation muscle activity (i.e., -25 – 0 ms). Unlike the case of the behavioral task during microneurography, the participants received no prior training in the main behavioral task with the robot. As the situation of interacting with a robot that perturbs one’s hand on every trial is also less than completely naturalistic, the initial five repetitions of each trial type were considered to be ‘familiarity’ trials and were excluded from analyses; excluding a number of initial trials is a common approach in similar robot-based sensorimotor control studies, e.g.,^56^. In experiment ‘2’, three preparatory delays were used (.25, .75 and 1.25 sec), reflecting the middle of each of the three epochs used for analyses in experiment ‘1’ (Fig. 2b). As expected from the afferent findings (Fig. 2c), visual inspection on EMG signals confirmed that a similar suppression of spinal SR occurred for the two longer delays (e.g., Fig. 5c-e represents trials were the delay was 0.75). The data were therefore collapsed across the two delays, to represent one ‘long’ delay condition (Fig. 6a-h). The relevant data used in statistical analyses for each participant were generated by first creating averages (medians) of EMG signals across repetitions of a relevant trial type that involved stretch of the particular muscle (i.e., EMG signals during muscle shortening were not analyzed in the current study as we were interested in stretch reflex responses). The average value within the epoch of interest was then taken, producing a single data-point per muscle and trial type. To simplify analyses (i.e., concentrate on the main manipulation of interest while accounting for known effects of e.g., muscle loading), for each individual muscle, EMG data of a particular load and/or delay were contrasted in terms of the target goal, generating a single data point that was ultimately used for statistical analyses as part of a single-sample t-test (see e.g., Fig. 6d, 6h and 7).

All statistical comparisons were two-tailed, and the overall baseline statistical significance level was 0.05. Tukey’s HSD test was used for any post-hoc analyses. No statistical methods were used for pre-determining sample sizes but the sizes used are similar to those reported in previous studies. Data normality was confirmed using the Shapiro-Wilks test for samples with <50 data-points and Lilliefors test for larger samples. Statistical tests were performed using either MATLAB® (MathWorks, Natick, MA, USA) or STATISTICA® (StatSoft Inc, USA).

## Acknowledgements

This work was supported by grants awarded to M.D. by the Kempe Foundation, the local Foundation for Medical Research (“Insamlingsstiftelsen”) and the Swedish Research Council (project 2016-02237). The funders had no role in study design, data collection and analysis, decision to publish or preparation of the manuscript.

## Author contributions

M.D. conceptualized and designed the study, M.D. collected the neural data, M.D. and S.P. analyzed the data, interpreted the results and wrote the manuscript.

---

**Supplementary Figure 1:**
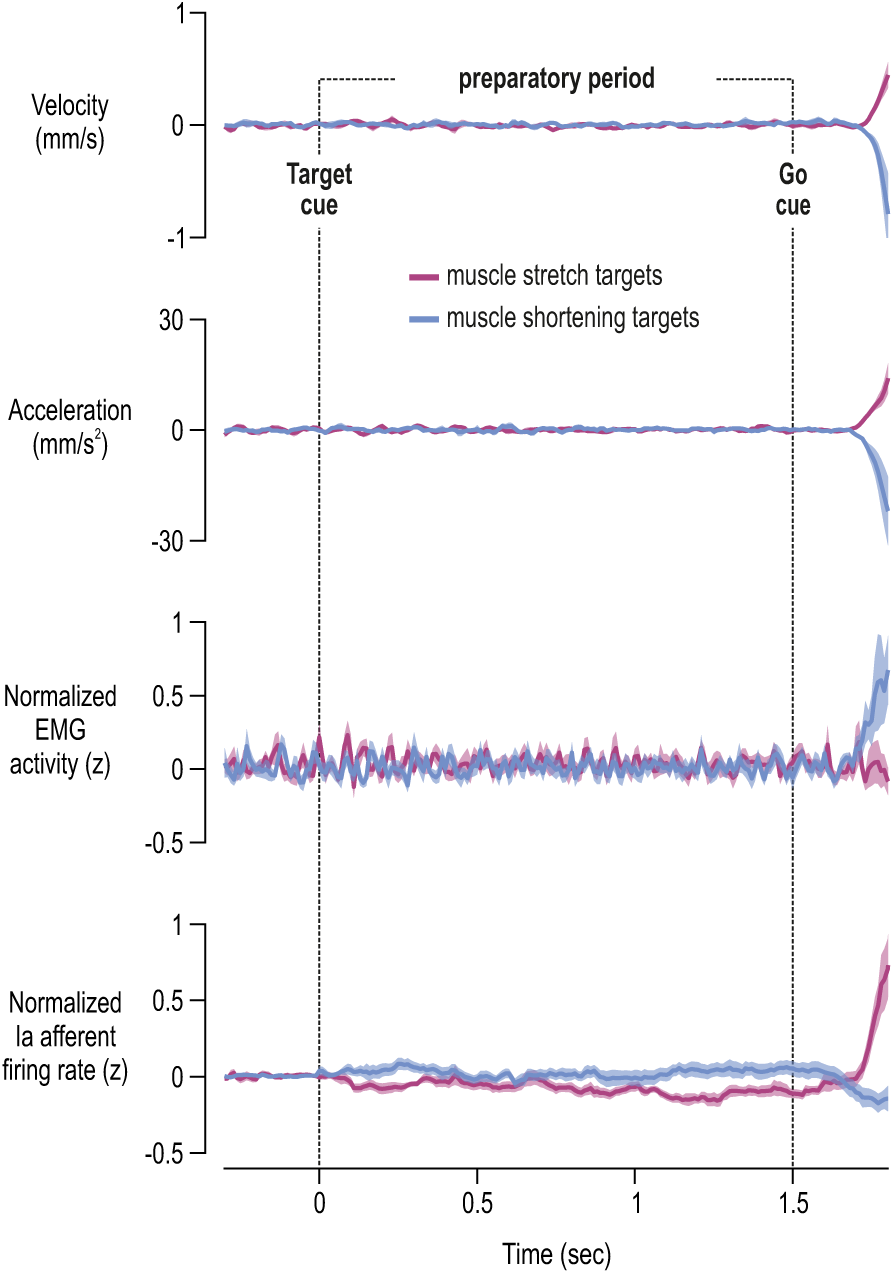
Population signals before, during and after movement preparation. Mean stretch velocity, acceleration, EMG and spindle type Ia signals across all recorded spindle-bearing muscles. The traces are aligned to onset of the target cue (time ‘0’) as per Figure 2b. Purple and blue traces represent targets associated with stretch and shortening of the spindle-bearing muscle, respectively. Shading represents ±1 s.e.m. Here, signals are also shown for the short period (0.3 sec) following onset of the Go signal where reaching movement begun to occur.

**Supplementary Fig. 2:**
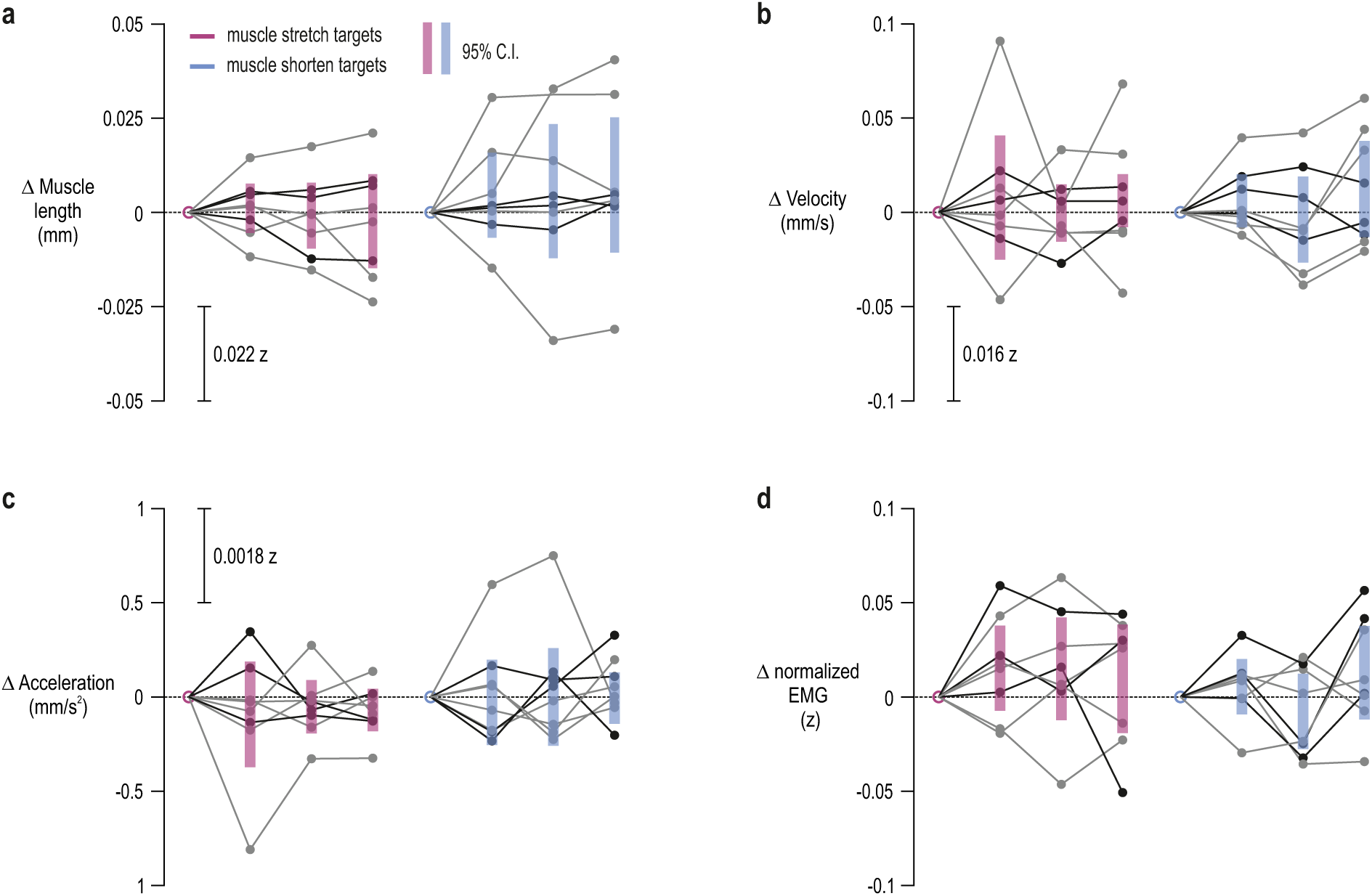
Very small deviations in kinematic signals and variability in EMG during preparation are unrelated to spindle tuning. **a-d**, Spindle-bearing muscle length, velocity, acceleration and EMG, respectively, corresponding to the afferent data presented in Figure 2c. Thin grey lines represent data from individual wrist extensor muscles and thin black lines represent data from digit extensors. The shaded bars represent 95% confidence intervals. The same color scheme is used throughout. As expected, deviations in these variables were minor and, importantly, none of the groups systematically differed from baseline, and no trends similar to those observed in Ia firing were seen (i.e., purple epoch ‘3’ < epoch ‘1’; Fig. 2c). Scales of normalized values (z) are also shown, reinforcing that deviations in these variables during preparation were very small compared to the changes observed across the full duration of the delayed-reach task (see also Fig. 3b-c).

**Supplementary Figure 3:**
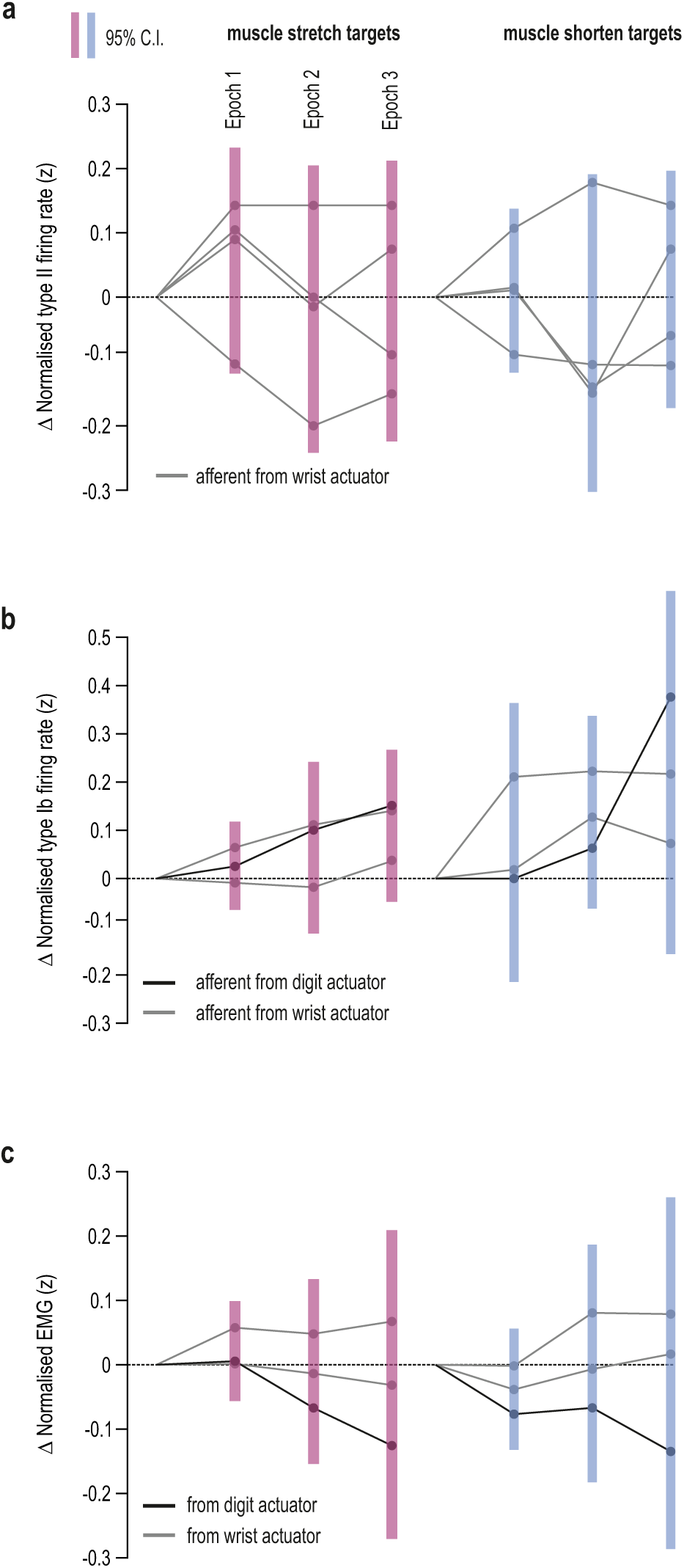
Type II and type Ib responses during movement preparation. **a** As Figure 2c but representing secondary muscle spindle afferents (‘type II’). **b** Same format as ‘a’ but representing afferent activity from Golgi tendon organ afferents (‘type Ib’). **c** Same format as ‘b’ but representing Golgi-bearing muscle EMG.

**Supplementary Figure 4:**
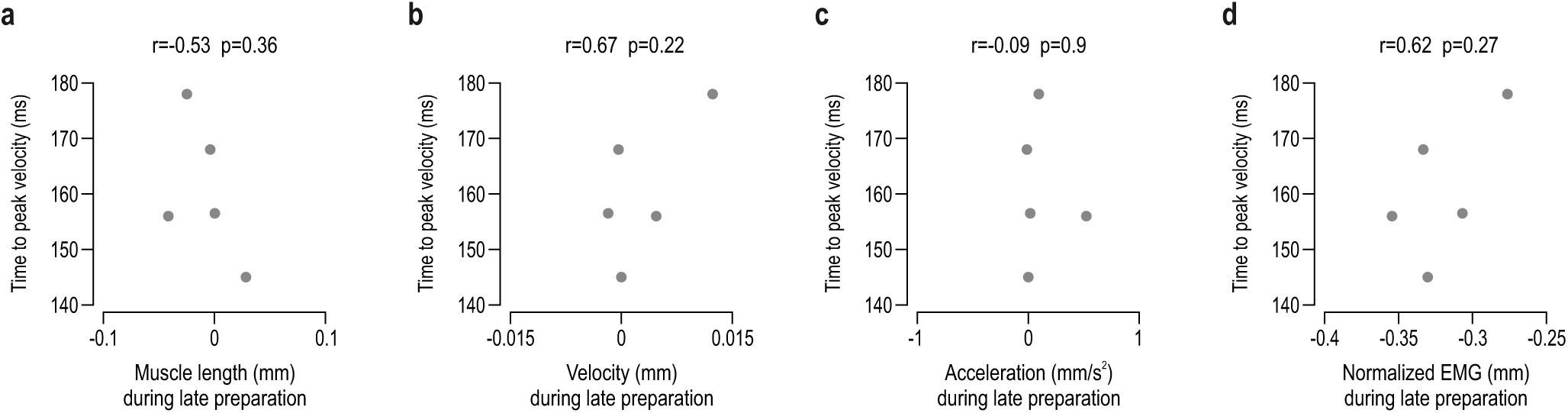
Kinematic signals and EMG at late movement preparation do not predict time to peak velocity. As Figure 4c but horizontal axes pertain to spindle-bearing muscle length (**a**), velocity (**b**), acceleration (**c**), and EMG (**d**). There was no significant relationship between any of these variables and time to peak velocity during reaching.

**Supplementary Figure 5:**
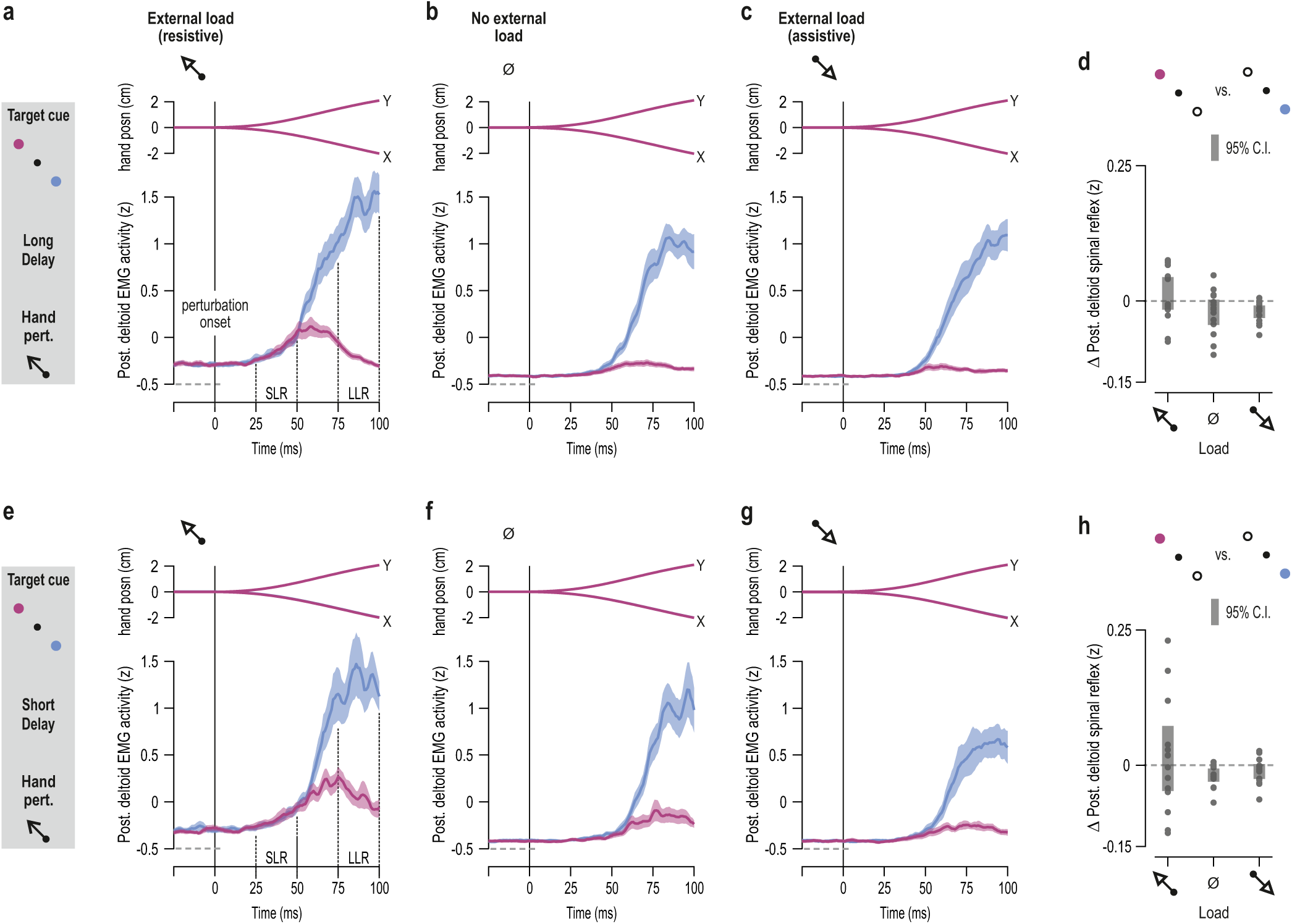
Equivalent goal- and delay-dependent effects on stretch reflex gains of the posterior deltoid. **a-c** Mean hand position (posn.) and mean rectified posterior deltoid EMG activity across participants when an external (pre-)load was applied in the direction of posterior deltoid stretch (‘a’), when there was no external load (‘b’), and when the load was applied in the direction of posterior deltoid shortening (‘c’). Shading represents ±1 s.e.m. As the schematic on the far left indicates, the data here represent trials where the preparatory delay was relatively long and the subsequent perturbation stretched the posterior deltoid. SLR denotes the epoch associated with the spinal stretch reflex and LLR the epoch associated with the long-latency stretch reflex (for LLR analyses see Fig. 7). **d** There was a consistent pattern of SLR modulation, equivalent to that observed for the pectoralis (i.e., Fig. 6d). **e-h** As top row of panels but representing trials where the preparatory delay was relatively short (250 ms).

**Supplementary Figure 6:**
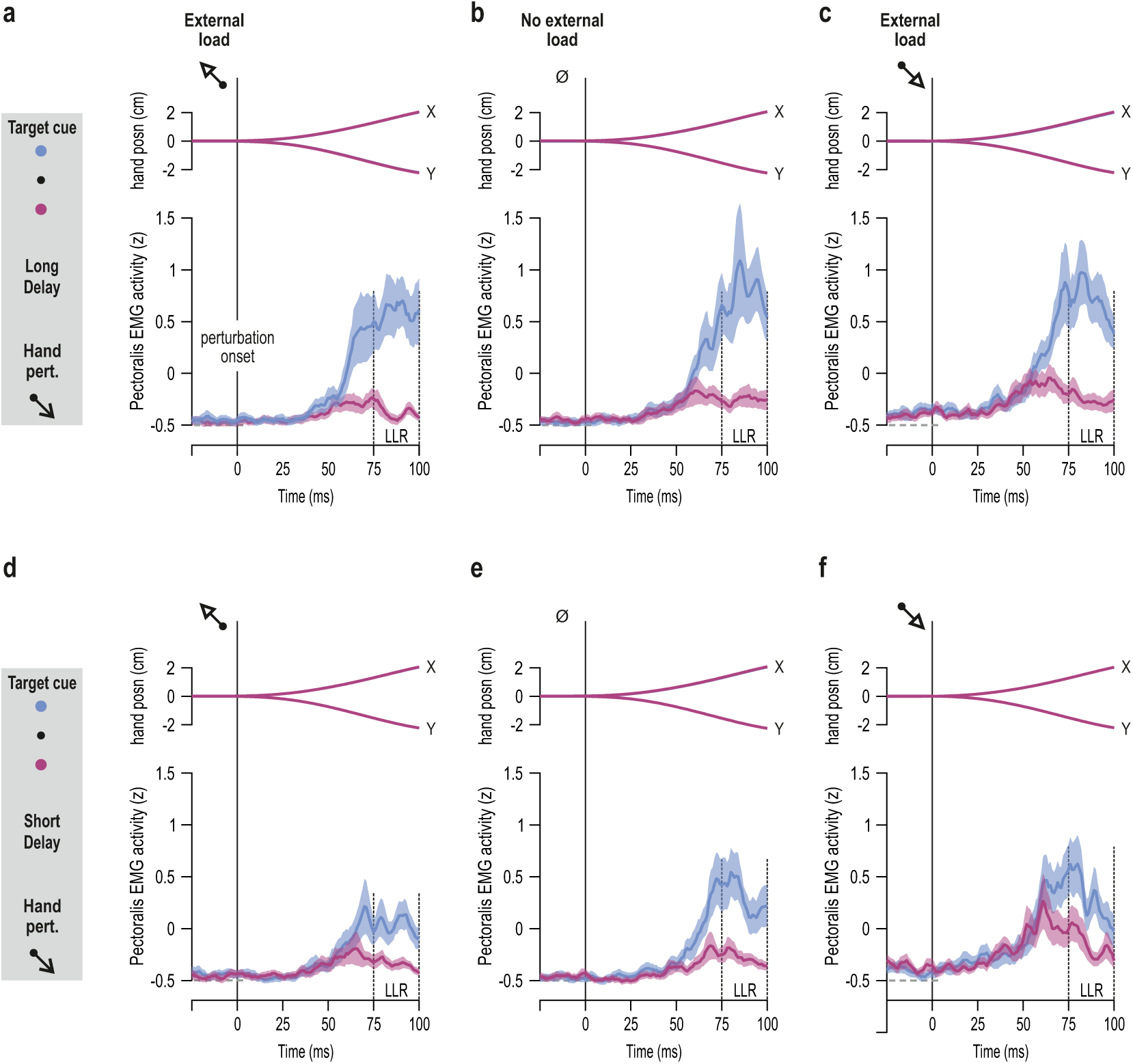
Goal- and delay-dependent modulation of LLR gains of the triceps muscle. As Figure 6 but pertaining to triceps lateralis when the targets required reaching along the vertical axis of the workspace. **a-c** Mean hand position (posn.) and mean rectified triceps laleralis EMG activity across participants (N=12) when the external load was applied in the direction of triceps shortening (‘a’), when there was no external load (‘b’), and when the triceps was externally loaded i.e., the load was applied in the direction of triceps stretch (‘c’). Shading represents ±1 s.e.m. As the schematic on the far left indicates, the data represent trials where the preparatory delay was relatively long (1.2 sec) and the subsequent perturbation stretched the triceps. LLR denotes the epoch associated with the long-latency or ‘R3’ stretch reflex response. **d-f** As top row of panels but representing trials where the preparatory delay was relatively short (0.2 sec).

